# BCG vaccination mitigates tau pathology and restores cognitive function in PS19 mice

**DOI:** 10.64898/2026.05.12.724591

**Authors:** Somnath Shee, Meixiang Huang, Meghraj Singh Baghel, Yuxin Zheng, Shichun Lun, Santosh K. Yadav, Nirbhay N. Yadav, Carlos E. Ruiz-Gonzalez, Sandeep Tyagi, Eric Nuermberger, Sanjay K. Jain, Zaver M. Bhujwalla, Barbara S. Slusher, Philip C. Wong, William Bishai

## Abstract

Retrospective studies in patients with non-muscle invasive bladder cancer (NMIBC) have reported a significant reduction in Alzheimer’s disease (AD) incidence (12–78%) among Bacillus Calmette-Guérin (BCG) recipients versus controls. To investigate the underlying mechanisms, we evaluated BCG in the PS19 mouse model of tauopathy. We found that BCG administration reduced hippocampal phospho–tau and microgliosis while preserving neuronal markers. *In vivo* volumetric T2-MRI demonstrated attenuation of brain atrophy accompanied by increased glutamate–weighted CEST–MRI signals. Functionally, BCG-treated mice showed improved performance in the novel object recognition test (NORT), as well as improved body-weight maintenance and survival. Transcriptomic profiling of the hippocampus revealed near complete normalization of the PS19 disease-associated gene expression signature towards that of healthy controls. Flow cytometric profiling of brain myeloid populations demonstrated a reduction in activated resident microglia, but total microglia cells remain elevated. Moreover, an increase of the co-stimulatory marker CD80 on the recruited peripheral myeloid cells ensues following BCG treatment. Consistent with this shift in myeloid state, primary brain myeloid cells from BCG-treated mice also exhibited enhanced phagocytosis of FITC–labeled tau fibrils and increased lactate production. Together, these findings indicate that BCG induces systemic and CNS myeloid cell reprogramming that limits neuroinflammation, enhances tau clearance, and rescues cognitive and neurodegenerative phenotypes in a tauopathy model. BCG is a safe, readily available therapy that merits consideration as a preventive agent against dementia.

**One sentence summary:** BCG therapy prevents tauopathy in PS19 mouse model.

## Introduction

There is a growing appreciation that vaccines may have beneficial off-target effects such as reducing the risk of dementia. Recent, well publicized studies of shingles vaccines have revealed compelling evidence that immunization against varicella-zoster virus may reduce the incidence of dementia. Capitalizing on age-based eligibility for zoster vaccination in Wales, it was shown that receipt of the live-attenuated zoster vaccine was associated with a 20% reduction in incident dementia over seven years. (1) Subsequent studies confirmed this same observation with the recombinant zoster vaccine. (2,3) And it has also been shown that shingles vaccination may influence multiple stages of the dementia disease course. (4)

While less publicized, there is also considerable evidence that Bacillus Calmette Guérin or BCG, the live attenuated vaccine to prevent tuberculosis (TB) and treat bladder cancer, may also have dementia-preventive properties. To date, six retrospective studies among NMIBC patients have each found a statistically significant reduction in the incidence of dementia in BCG-recipients compared to controls who did not receive BCG. (fig. S1) (5–10) Across these six studies the reductions ranged from 12-78%, and the total number of patients monitored was >85,000 with mean follow-up times of 3 to 15 years. Three meta-analyses confirmed the significance of the retrospective bladder cancer studies. (11–13) In addition, a 49-patient prospective trial of cognitively unimpaired older adults found that two intradermal doses of BCG over four weeks improved an objective blood biomarker measured nine months later. (14) BCG and other vaccines have been shown to stimulate protection against heterologous antigens by a process known as trained immunity. Upon exposure to a first antigen such as BCG, innate immune cells, typically myeloid cells, have been shown to undergo epigenetic and metabolomic reprogramming such that upon exposure to a second, unrelated antigen their immune response is greater than it was to the first. (15–17) Indeed, trained immunity has been postulated as a mechanism by which BCG may protect against dementia. In this regard, it is noteworthy that a recent study of chemically-induced demyelination in aged C57BL/6 mice found that intravenous BCG administered one month prior to the injury dramatically accelerated healing via a mechanism involving H3K27ac and H3K4me3 epigenetic activation marks in the microglial/peripheral myeloid cells. (18)

To directly test BCG as a preventative therapy for AD, we employed the PS19 (P301S) mouse model of tauopathy. (19–21) This transgenic mouse expresses the P301S mutant form of the human microtubule-associated protein tau (MAPT; 1N4R) on chromosome 3 under the control of the mouse *Prnp* promoter. (19) The overexpression of P301S tau is associated with AD-relevant pathological and functional features, including neuronal loss, microgliosis, neurofibrillary tangles (NFT) deposition, and cognitive impairment. (20,21) Moreover, aberrant microglial activation is both a driver and consequence of tau pathology in this model. As phospho-tau (p-tau) accumulates in the PS19 mouse brains, microglial cells are seen to transition from a homeostatic, surveillant state to an activated phenotype that amplifies proinflammatory cytokine signaling, complement cascades, and contributes to synaptic loss. (19,22–25) Because microglia and peripheral monocytes are central to NFT clearance and maintenance of neuronal homeostasis, (26,27) we hypothesized that BCG-mediated training of myeloid cells may rebalance their state from the pathologic chronically activated state to an efficient phagocytic and homeostatic state, and that such transitions may delay neuronal loss in the model.

In this study, we show that subcutaneous BCG-administration mitigates hallmark features of tauopathy in PS19 mice. BCG reduced hippocampal phospho-tau burden and microgliosis while preserving neuronal markers. BCG-treated PS19 mice preserved hippocampal and ventricular volume, displayed improved survival, body weight and recognition-memory performance. This protective effect correlated with a BCG-driven myeloid cell reprogramming towards enhanced glycolysis and phagocytic capacity, higher co-stimulatory CD80-expression on peripheral myeloid cells with concomitant dampening of microglial activation and proinflammatory cytokine signaling. In sum, our data show that BCG significantly reduces tau pathology in the PS19 model and provides a mechanistic basis for BCG-mediated AD-mitigation.

## Results

### Subcutaneous BCG-administration reduces hippocampal p-tau deposition and microgliosis, while preserving neuronal integrity

The B6;C3-Tg(Prnp-MAPT*P301S)PS19Vle/J or PS19 mouse model has been reported to display p-tau seeding at 3 months of age, neuronal loss and microglial activation at 6-months of age, and prominent NFT deposition at 6-9 months of age. (19) The median survival time of PS19 mice is 10-12 months indicating aggressive tauopathy. (19,28) To examine whether BCG therapy may attenuate disease progression in this model, hemizygous PS19 mice were either injected with phosphate buffered saline (1X PBS; sham) or with BCG (2×10^6^ colony-forming units (CFU)) subcutaneously in the dorsal neck (scruff) region. Age-matched, non-PS19 littermate mice (Non carriers) were sham-treated and represent untreated, age-matched healthy controls (AMC UT). Three monthly BCG doses were administered starting at 3-months of age, followed by a no-treatment phase of 3-4 months (Fig. 1A). The hippocampus and prefrontal cortex are the major regions affected by the tauopathy in the PS19 model. (19) Therefore, mice were sacrificed at 9-10 months of age, and disease in the hippocampus and cortex was assessed by immunohistochemistry (IHC) with antibodies directed against pSer202/pThr205 phospho-tau (AT8 anti-p-tau antibody), Iba1 (microglial activation marker), GFAP (astrocyte activation marker), and NeuN (neuronal integrity marker) (fig. S2, Fig. 1). While p-tau depositions in the hippocampus were higher in hemizygous PS19 mice relative to age-matched controls (48-fold increase; *p*<0.0001), BCG-treatment significantly reduced the percent AT8 stained area (5-fold reduction relative to that of sham-treated PS19 animals; *p*=0.0002) (Fig. 1B). In addition, BCG treatment also significantly reduced microglial activation (an indicator of neuroinflammation) as measured by the percent Iba1+ stained area in the hippocampus CA1 region (4-fold reduction relative to that of sham-treated PS19 mice; *p*=0.002), and restored homeostatic-state of microglia (Fig. 1C, fig. S3A). Importantly, BCG also served to prevent neuronal death as demonstrated by the increased percent NeuN+ stained area in the hippocampal CA1 region and cortex, compared with sham-treated PS19 mice (Fig. 1E, fig. S3B-C). Interestingly, BCG therapy did not reduce astrocyte activation as measured by GFAP immunoreactivity which also occurs in the hippocampus of PS19 mice (fig. S3D-E). Taken together, the IHC data show that BCG administration ameliorated tauopathy disease progression, neuroinflammation, and associated neuronal death in PS19 mice.

**Fig. 1.**
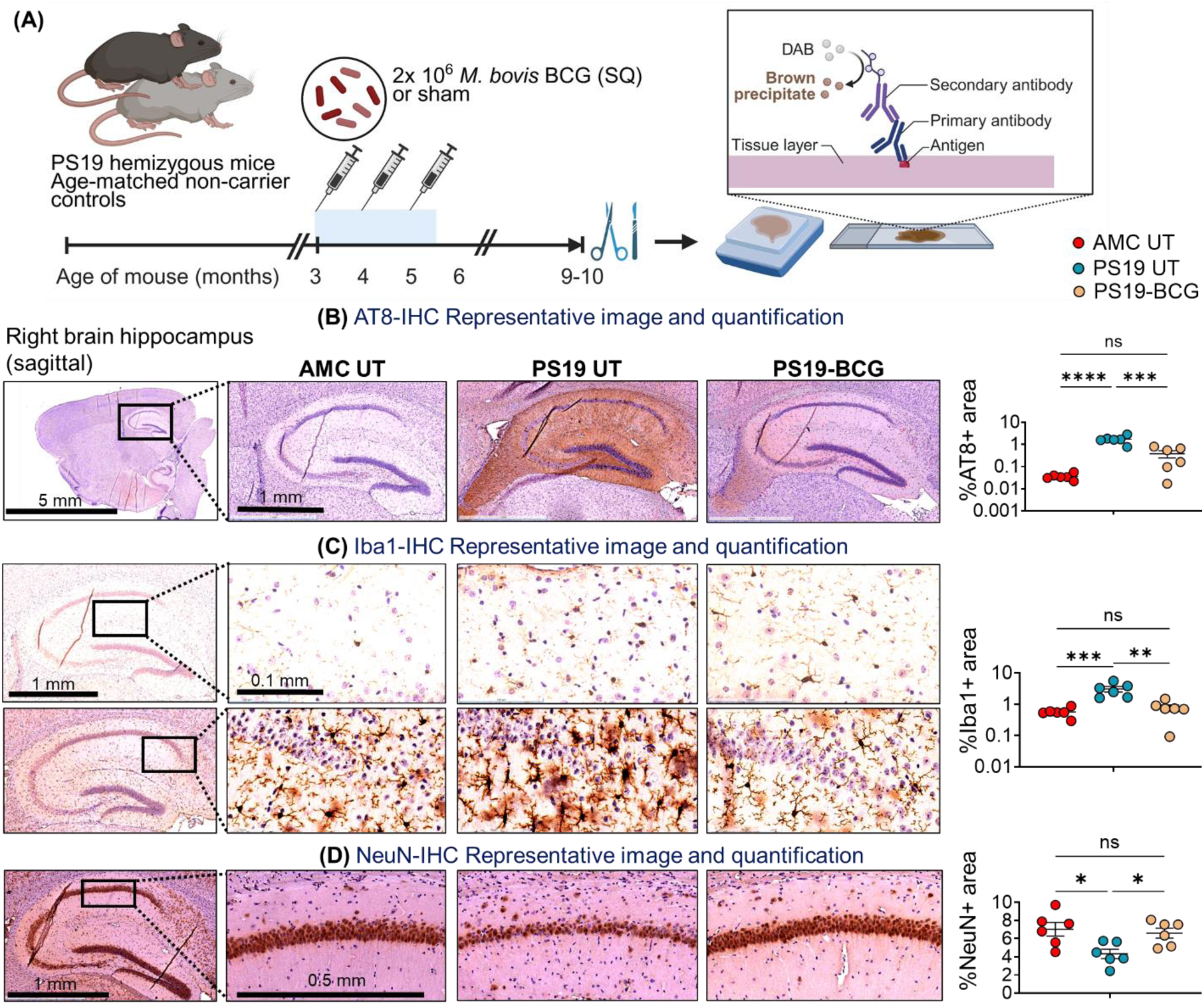
*M. bovis* BCG treatment ameliorates p-tau accumulation, microgliosis, and neurodegeneration in PS19 mice. **(A)** Schematic of the experiment. **(B)** %AT8+ area in the hippocampus of age-matched controls (AMC untreated (UT)), Sham (PBS) or BCG treated PS19 mice, as determined by ImageJ analysis. (n=6/group) **(C)** %Iba1+ area in the hippocampus-CA1 region of AMC UT, Sham (PBS) or BCG treated PS19 mice, as determined by ImageJ analysis. (n=6/group) **(D)** %NeuN+ area in the hippocampus-CA1 region of AMC UT, Sham (PBS) or BCG treated PS19 mice, as determined by ImageJ analysis. (n=6/group). The data are means ± SEM, and representative of two independent experiments. For each mouse brain sample, IHC was done in three planes (10 µm thick sections) for all the antibodies. Each data point represents a mouse. Statistical analysis was calculated by one-way ANOVA with Tukey’s multiple comparisons test. (*p*>0.05: ns, *p*<0.01: **, *p*<0.001: ***, *p*<0.0001: ****)

### BCG treatment prevents weight loss and extends survival of PS19 mice

Microglial activation and neuronal cell loss are closely associated with morbidity such as reduced mobility and weight loss, and these lead to early mortality in the PS19 model. (19) By 8-9 months of age, the body weights of hemizygous PS19 mice show significant reductions relative to the age-matched non-carrier controls, and the difference increases with age. (28) Moreover, the PS19-tau pathology is more severe in males than females, and hence males lose weight faster and have shorter survival times than females. (28) We therefore tested whether BCG treatment influences body weight and survival in two independent mouse cohorts (Fig. 2A). In cohort 1 (25 male and 30 female PS19 mice), we administered PBS (sham) or 2×10^6^ CFU of BCG to hemizygous PS19 mice in 3 monthly doses. As expected, sham-treated PS19 mice displayed reduced survival (*p*=0.06) and failed to maintain body weight compared to age-matched controls through 9 months of age (Fig. 2B, 2C). In contrast, BCG treatment reversed the accelerated mortality phenotype (*p*=ns compared to age-matched controls, Fig. 2B) and led to a gain of body weight at 5-6 (8%) and 7-8 (15%; *p*=0.02) months of age relative to the sham-treated PS19 mice (Fig. 2C).

**Fig. 2.**
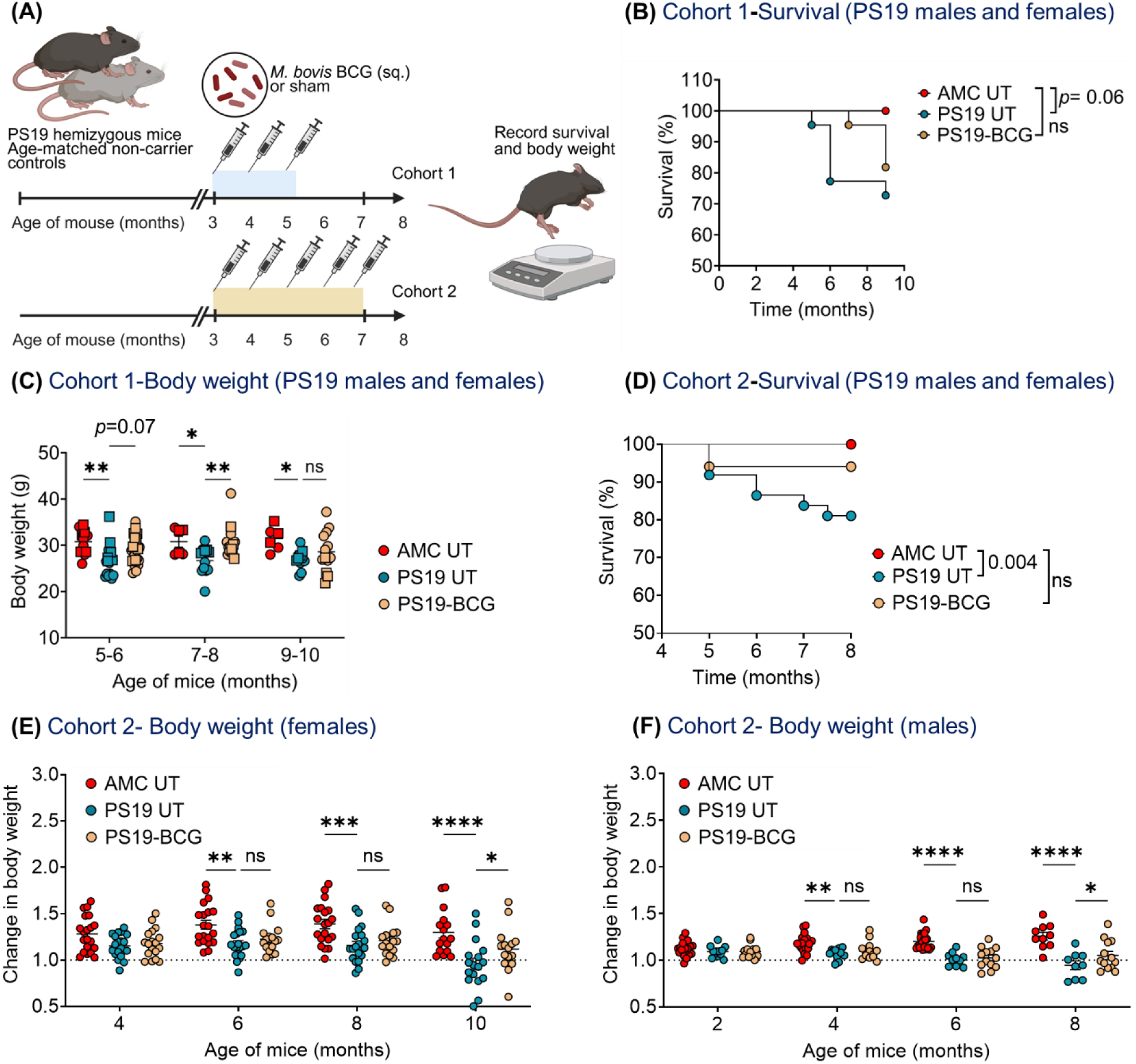
*M. bovis* BCG treatment is safe and improves body weight and survival of PS19 mice. **(A)** Schematic of the experiment. Cohort 1 data: **(B)** Kaplan-Meir survival curve of age-matched controls (n=11), sham-treated (n=22) and BCG-treated (n=22) PS19 males and females’ mice. Indicated *p* values are measured by log-rank (Mantel-Cox) test. **(C)** Changes in body weight of the surviving mice. In Cohort 2, PS19 mice were treated with 5 monthly doses of BCG. **(D)** Kaplan-Meir survival curve of age-matched controls (n=40), sham-treated (n=37) and BCG-treated (n=34) PS19 males and females’ mice. Indicated *p* values are measured by log-rank (Mantel-Cox) test. Changes in body weight of the surviving **(E)** females and **(F)** males from Cohort 2. The data are means ± SEM. Each data point represents a mouse. Statistical analysis was calculated by two-way ANOVA with Dunnett’s multiple comparisons test. (*p*>0.05: ns, *p*<0.01: **, *p*<0.001: ***, *p*<0.0001: ****)

Since BCG injections are safe both in newborn humans and mice, (17) we increased the dosage of BCG in the cohort 2 (67 male and 77 female PS19 mice) to determine whether treatment outcomes are dose-dependent. We administered PBS (sham) or 10×10^6^ CFU of BCG to hemizygous PS19 mice in 5 monthly doses. Similar to cohort 1, BCG significantly improved survival of PS19 mice (6% died; *p*=ns relative to AMC UT) (Fig. 2D). Moreover, BCG-treated PS19 mice displayed body weight gain (10-20%) relative to the sham-treated PS19 mice at 10 (Fig. 2E; females) and 8 months of age (Fig. 2F; males). In sum, the magnitude of body-weight improvement was broadly similar across the two treatment regimens.

### BCG treatment prevents hippocampal atrophy and reduces ventricular dilatation

Tauopathy in PS19 mice follows a close trajectory to tauopathy syndromes seen in humans with the mice developing significant volume loss of the hippocampus, amygdala, hypothalamus and cortex, with associated ventricular dilatation by 9 months of age. (19,29,30) We examined whether BCG administration may prevent brain atrophy in PS19 mice using T2-weighted magnetic resonance imaging (T2-W-MRI). We focused on hippocampal atrophy and ventricular dilatation and used the animals in Cohort 1 and Cohort 2 (Fig. 2A) for these live-imaging studies. T2-W-MRI with 6–7-month-old PS19 mice from cohort 1 or age-matched controls did not reveal significant differences in the hippocampus volume (fig. S4A, S4B); although we observed a modest increase in ventricular volume in sham-treated 6–7-month-old PS19 mice that was reduced by BCG therapy, these changes were not statistically significant (fig. S4B). Interestingly, at this time point, we observed reduced total human tau protein levels (31) in serum from the BCG-treated PS19 mice (fig. S4C; *p*= 0.008). In cohort 2, we increased the BCG dose by subcutaneously administering PBS (sham) or BCG (10×10^6^ CFU) into hemizygous PS19 mice in 5 monthly doses. As expected, we observed distinct ventricular dilatation in 9 months old PS19 females, as shown as high-intensity areas in the T2-W axial sections (Fig. 3A-D). In addition, hippocampal atrophy was evident both in the axial sections and in 3D reconstructions in sham-treated PS19 females (Fig. 3C, 3D). Volumetric analysis by ITK-SNAP software (32) indicated a 21% loss of hippocampal volume and a 63% increase in ventricular volume in sham-treated PS19 females, both of which were reversed by BCG treatment (Fig. 3B). Due to the early onset of paresis in sham-treated PS19 males, we evaluated atrophy in males at 8 months of age (fig. S4D-F). Interestingly, while we did not see changes in hippocampus volume at this time-point, we see ventricular dilatation in untreated PS19 males which was significantly reduced in PS19 males treated with BCG (fig. S4F).

**Fig. 3.**
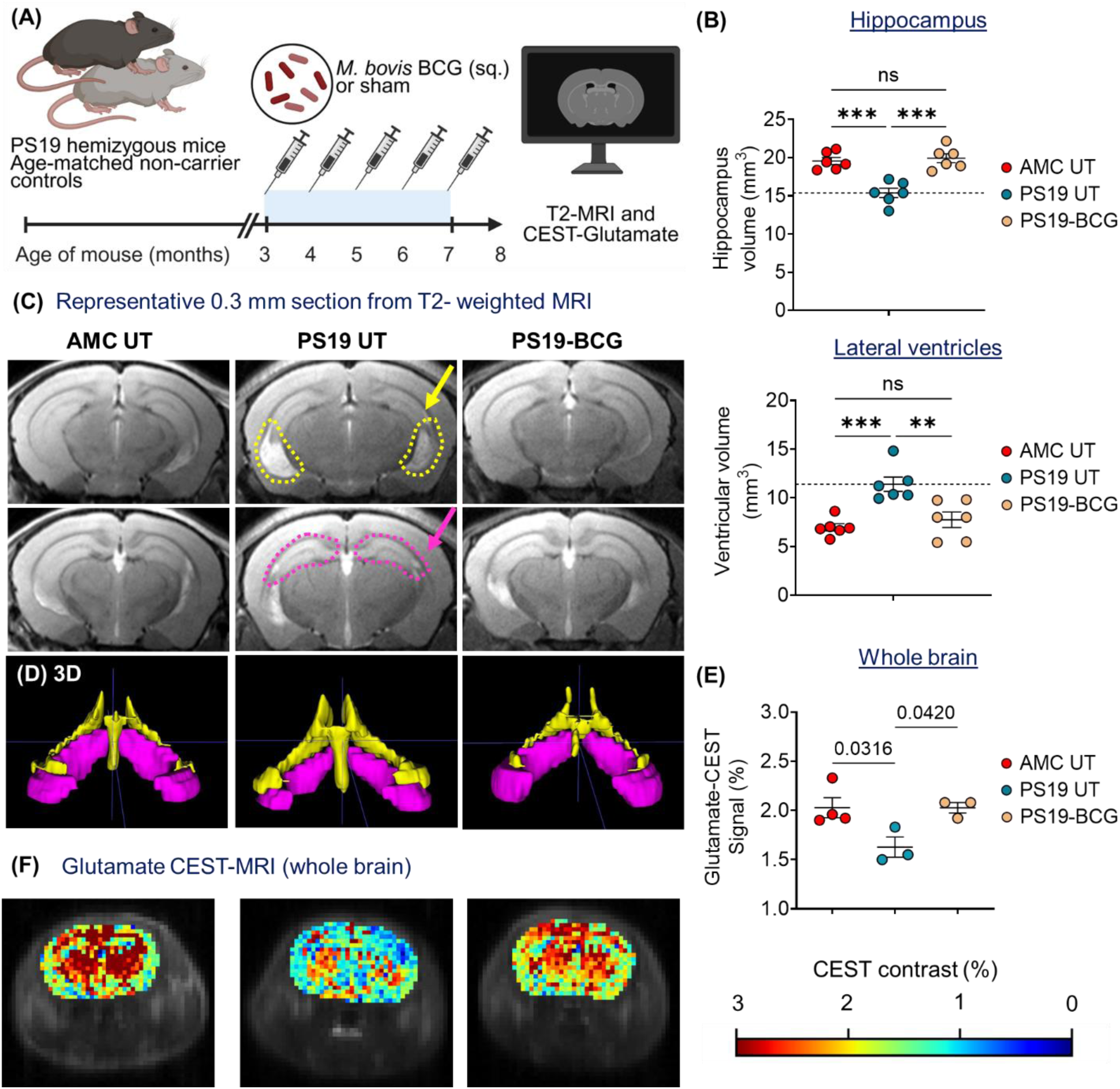
*M. bovis* BCG treatment prevents atrophy in the brain of PS19 mice. **(A)** Schematic of the experiment. **(B)** Quantitative analysis of the hippocampus and lateral ventricle volume as determined by using ITK-SNAP software. (n=6/group) Statistical analysis was calculated by one-way ANOVA with Tukey’s multiple comparisons test. **(C)** Representative 0.3 mm section of *in vivo* T2-weighted MRI illustrating significant atrophy of hippocampus (encircled with magenta) and ventricular dilatation (encircled with yellow) in 8–9-month-old PS19 mice. Yellow and pink arrow indicate ventricular enlargement and hippocampus atrophy respectively, in sham-treated PS19 mice. **(D)** Representative 3D rendering of hippocampus and lateral ventricles, as done by ITK-SNAP software. **(E, F)** Representative image of glutamate-weighted chemical exchange saturation transfer (CEST-MRI) maps of the brain. (n=3 mice/group), and quantification. The data are means ± SEM, and representative of two independent biological experiments. Each data point represents a mouse. Statistical analysis was calculated by one-way ANOVA with Dunnett’s multiple comparisons test. (*p*>0.05: ns, *p*<0.01: **, *p*<0.001: ***, *p*<0.0001: ****)

Recent studies have demonstrated that glutamate levels decrease as early as 7 months of age in live PS19 mice, and this change can be measured in-vivo by an MRI method called glutamate-weighted chemical exchange saturation transfer (GluCEST-MRI) imaging. (33,34) This non-invasive imaging method revealed significant- 15-30% decreases in glutamate levels in the hippocampus and thalamus regions in PS19 mice. (33,34) Therefore, we measured glutamate levels in randomly selected mice from each treatment group by GluCEST-MRI. We found that the whole brain mean GluCEST signal was significantly lower in the sham-treated PS19 mice relative to age-matched controls and that BCG administration restored the GluCEST signal to that seen in healthy, untreated non-PS19 mice. (Both Cohort 1 (fig. S4G, H) and Cohort 2 (Fig. 3E, F)). Taken together, these non-invasive imaging data show that BCG treatment prevents tauopathy-driven brain atrophy and allows maintenance of normal glutamate levels in the PS19 mice.

### Subcutaneous administration of BCG alleviates cognitive defects

Given that BCG treatment reduces pathological tau burden, preserves neuronal integrity (NeuN), and dampens microglial activation (Iba1), and that these changes are accompanied by reduced brain atrophy by MRI, we next assessed whether these neurological benefits translate into improvements in cognitive performance. To evaluate the functional impact of BCG treatment on recognition memory, we performed behavioral testing using the novel object recognition test (NORT). (29,35) PS19 hemizygous mice and age-matched non-carrier (AMC) controls were assessed at 8 months of age, following 5 months of vehicle or BCG treatment (Fig. 4A). Locomotor activity was first evaluated using the open field test. Total beam breaks over 30 minutes did not differ significantly among groups (Fig. 4B), indicating preserved motor function and comparable exploratory activity across cohorts. Recognition memory was then assessed using the NORT. Untreated PS19 mice showed a significant reduction in recognition index compared to AMC controls (*p*=0.007), consistent with impaired novel object recognition memory (Fig. 4C). Importantly, BCG treatment significantly improved recognition performance of PS19 mice, restoring behaviors to levels approaching those of healthy AMC controls (*p*< 0.05; Fig. 4C). Similarly, the discrimination index was reduced in untreated PS19 mice compared to AMC controls (*p*=0.006), and was significantly rescued by BCG treatment (*p*< 0.05; Fig. 4D). Together these data demonstrate that BCG-mediated immunomodulation confers robust behavioral and cognitive improvements in a tauopathy mouse model without affecting locomotor function.

**Fig. 4.**
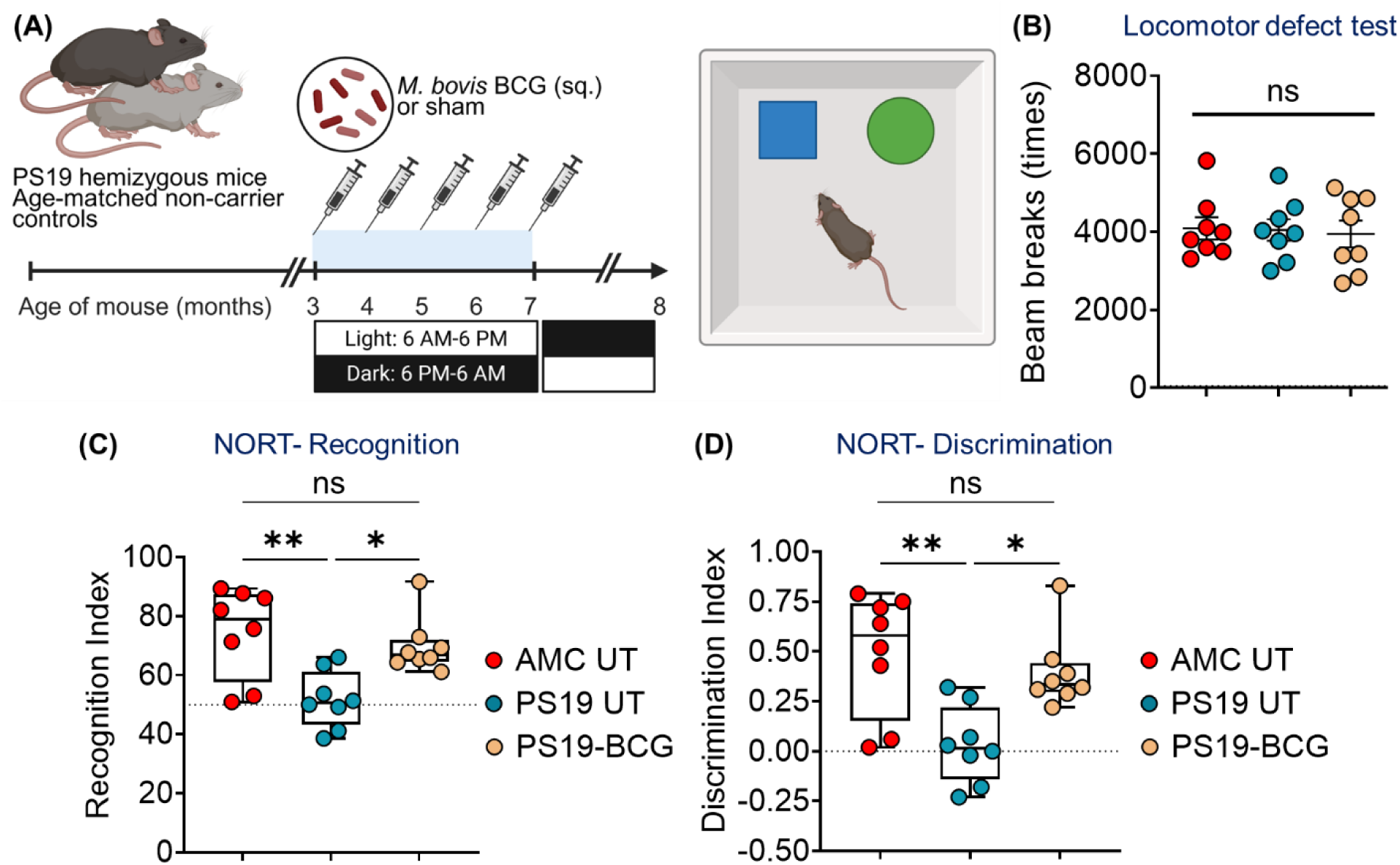
Subcutaneous *M. bovis* BCG treatment improves cognitive function in PS19 mice. **(A)** Schematic of the experiment. **(B)** Total beam breaks in the Open Field Test. **(C)** Recognition index and **(D)** discrimination index in the Novel Object Recognition Test. The data are means ± SEM, and representative of two independent experiments. Each data point represents a mouse. Statistical analysis was calculated by one-way ANOVA with Dunnett’s multiple comparisons test (B) and Kruskal-Wallis test with Dunn’s multiple comparison test (C, D). (*p*>0.05: ns, *p*<0.01: **, *p*<0.001: ***, *p*<0.0001: ****)

### BCG treatment restores a homeostatic transcriptional landscape and reduces proinflammatory signaling

To decipher cellular mechanisms that may explain the BCG-mediated protective effects we investigated the transcriptomics of hippocampal tissue in our studies; we selected the hippocampus because it is the region most heavily affected by the tauopathy process in PS19 mice. (19,30) Sham- and BCG-treated PS19 mice as well as untreated, age-matched, non-carrier controls who received monthly treatments at 3, 4, and 5 months of age were euthanized at age ∼9.5 months (Fig. 5A). Excised hippocampus tissue was used to isolate total RNA, and bulk RNA-sequencing was performed. As may be seen in Fig. 5B, untreated PS19 mice (3 middle columns) showed a dramatically different transcriptional pattern than untreated, healthy, non-PS19 AMC mice (3 left columns). Remarkably, subcutaneous BCG treatment of PS19 mice 4.5 months prior fully restored the transcriptional pattern (3 right columns) to essentially the same as that seen in untreated, healthy, non-PS19 AMC mice (3 left columns). Indeed, the two transcriptomes were so similar that in comparing BCG-treated PS19 mice to the AMC mice, there were only 9 differentially expressed genes (DEGs; *p.adj.*< 0.05, log_2_fold change >1 or <-1): three upregulated genes and six downregulated genes. (Supplementary file S1)

**Fig. 5.**
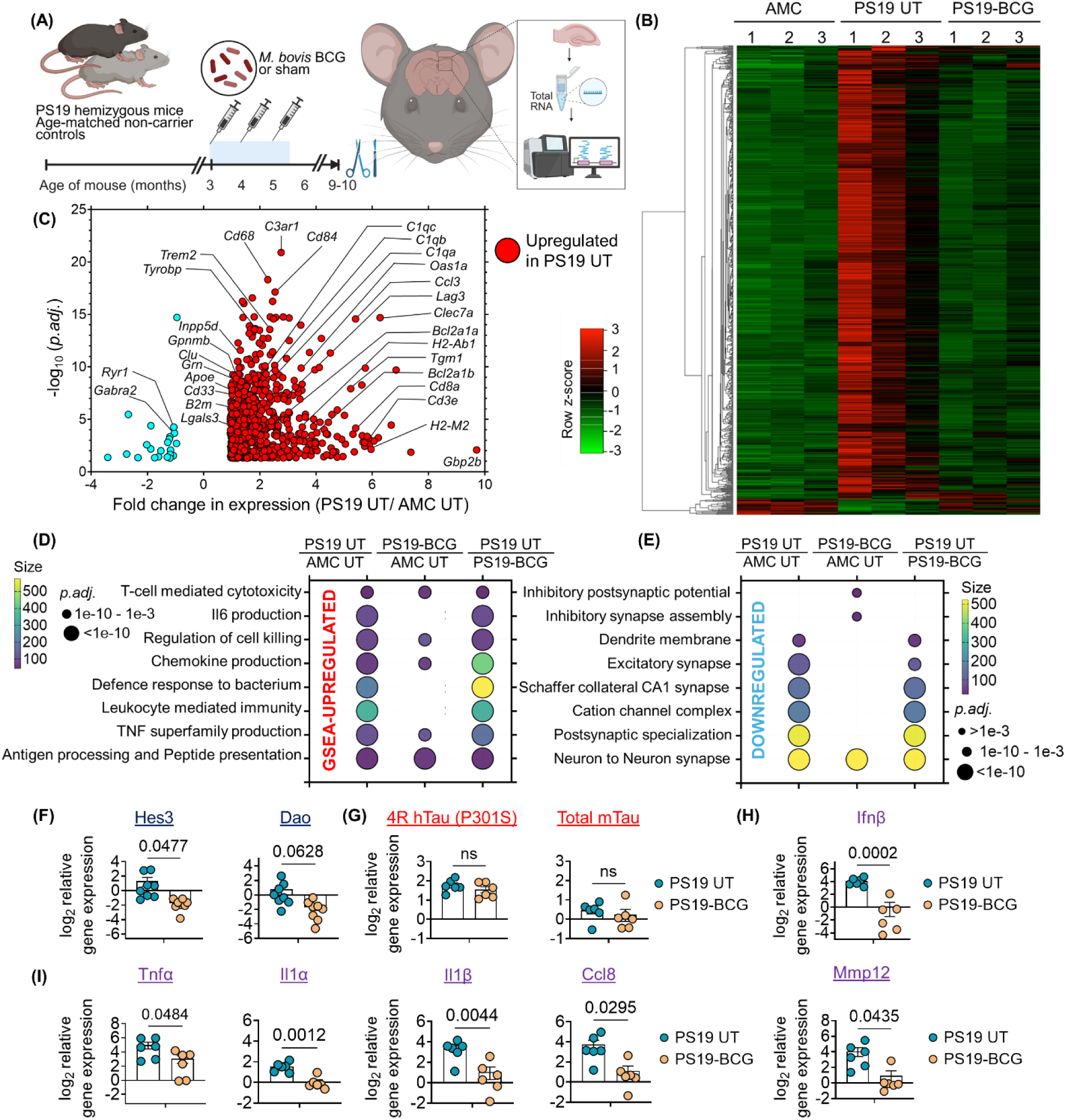
*M. bovis* BCG treatment restores homeostatic/resting transcriptional landscape in the hippocampus of PS19 mice. **(A)** Schematic of the experiment. Bulk RNA-Sequencing of hippocampus tissue (n=3 mice/group). **(B)** Heatmap showing transcripts per million (TPM) values of all 716 differentially expressed genes (DEGs; *p.adj.*< 0.05, log_2_fold change >1 or <-1) in the hippocampus from sham-treated PS19 mice relative to similarly treated age-matched non carrier controls. Red and green color indicates upregulated and downregulated genes in sham-treated PS19 mice, respectively. BCG treatment restores the transcriptional landscape of PS19 mice to that of age-matched controls. **(C)** Volcano plot showing major AD-associated genes to be upregulated (in red) in sham-treated PS19 mice relative to similarly treated age-matched non carrier controls. GSEA analysis revealed **(D)** topmost upregulated pathways, and **(E)** top downregulated pathways to be enriched under different comparisons, as indicated. **(F)** quantitative real-time PCR showing gene expression of **(F)** Hes3, Dao, **(G)** human tau (P301S), total endogenous mouse tau (m-Tau), **(H)** proinflammatory cytokines relative to age-matched controls in the hippocampus. Expression of targeted genes were normalized to housekeeping gene *Gapdh*. n= 6-8 mice/group. The data are means ± SEM, and representative of two independent experiments. Each data point represents a mouse. Statistical analysis between two groups was measured by unpaired two-tailed Student’s t test and *p* values are shown in the figure. (*p*>0.05: ns)

In contrast, the comparison of sham-treated PS19 mice to untreated age-matched, non-carrier control, revealed 716 DEGs: 692 upregulated and 24 downregulated genes (Fig. 5B, 5C, Supplementary file S1). In line with previously published data, upregulated genes in untreated PS19 mice included known risk-factor genes for AD (*Apoe, Trem2, Clu, Tyrobp, Grn, Cd33, Inpp5d, Oas1*). In addition to this, genes involved in complement cascades (*C1qa, C1qb, C1qc, C1rl, C3, C3ar1, C4b*), RIG-I-like receptor (RLR) signaling (*Ddx60, Ifih1, Ddx58, Dhx58, Zbp1, Isg15, Rsad2, Oas1b, Oas1g, Oas3, Oasl1*), senescence (36) (*Cdkn1a, Bcl2a1a, Bcl2a1b, Serpine 1, Plaur, Plau, Cxcl16, Lcp1, Csf2rb, Ang, Icam1*), cytokine–cytokine receptor interaction (*Il1b, Il18r1, Il18rap, Tgfb1, Cxcl5, Ccl3, Ccl5, Cxcl10, Cxcl13, Cntf, Tnfsf13b, Tnf, Adora3, Csf1r, Cxcr3, Cx3cr1, Il1rn*) were also upregulated relative to age-matched controls. Moreover, functional pathway analysis by GSEA software (Gene set enrichment analysis) (37,38) revealed “Antigen processing and peptide presentation” (Normalized enrichment score (NES)= 2.97), “Leukocyte mediated immunity” (NES= 2.87), “TNF superfamily cytokine production” (NES= 2.88), “chemokine production” (NES=2.84, *p.adj.*= 10^-15^), and “Il-6 production” (NES=2.76) to be among the top 20 pathways to be enriched in sham-treated PS19 mice, clearly indicating a heightened level of inflammation. (Fig. 5D). On the other hand, pathways that were significantly down-regulated in untreated PS19 mice relative to untreated AMC mice included “Neuron to neuron synapse” (NES=2.84), “Postsynaptic specialization” (NES=2.82), and pathways associated with neuronal homeostasis, consistent with ongoing neurodegeneration in the untreated PS19 hippocampi as seen by our previous NeuN IHC-staining (Fig. 5E). Taken together, the DEGs reflect p-tau-driven neurodegeneration and inflammation in the hippocampus.

Despite the remarkable similarity between the AMC and BCG-treated PS19 transcriptome, we next asked whether we could discern any differences between the two that might provide cues as to the BCG mechanism of action. GSEA functional pathway analysis revealed “Antigen processing and peptide presentation” (NES= 2.86) and “MHC Protein complex” (NES= 2.73) to be top-enriched suggesting continued innate immune cell activity in the BCG-treated PS19 mice. (Fig. 5D) However, pathways involved in “cytokine or chemokine production” were relatively less enriched (*p.adj.*= 10^-5^), suggesting a dampened proinflammatory milieu compared to untreated PS19 animals. Additionally, the downregulated transcriptome in BCG-treated PS19 mice displayed a distinct pattern when compared to age-matched controls; while we still observed “neuron to neuron synapse” to be significantly downregulated, inhibitory synapse-related gene-sets emerged among the topmost downregulated classes. These included “inhibitory postsynaptic potentials” (NES=-2), “regulation of presynaptic membrane potential” (NES=-2.02), and “inhibitory synapse assembly” (NES=-2.02; Leading edge= *Cbln1, Gabrb3, Srgap2, Gabra6, Cbln4, Gabra2, Gabrb2, Gabra1, Mdga1, Lgi2, Hapln4, Sema4a, Npas4, Gabrg2, Fgf13,* which represent genes involved in GABAergic inhibitory synapses and assembly of other inhibitory synapses) (Fig. 5E, Supplementary file S1). Taken together, the GSEA analysis suggests a compensatory model in which untreated PS19 hippocampus is dominated by a repression of excitatory synaptic plasticity programs, while BCG treatment drives the transcriptome toward a baseline-like state characterized by downregulation of inhibitory synaptic gene expression.

Lastly we observed that the expression of three genes was uniquely repressed in BCG-treated PS19 relative to sham-treated PS19, but were unchanged in AMC relative to sham-treated PS19: Hes3 (hes family bHLH transcription factor 3; log_2_foldchange= −5.3, *p.adj*=0.02), Dao (D-amino acid oxidase; log_2_foldchange= −2.7, *p.adj*=0.04) and Gm32113 (log_2_foldchange= −5.9, *p.adj*=0.005) (Fig. 5F, Supplementary File S1). It has been reported that reduced levels of the *Hes3* transcriptional repressor accelerates differentiation of progenitor cells into neurons, (39) suggesting that BCG-mediated Hes3 repression may aid in recovery from neuronal loss in the PS19 mice. Additionally, D-amino acid oxidase *Dao* mediates flavin adenine dinucleotide (FAD)-dependent oxidation of D-serine, D-alanine, D-proline, and D-leucine. (40–42) Since D-serine is a major agonist of N-methyl-D-aspartate (NMDA) receptor (glutamate-gated ion channels essential for synaptic plasticity), Dao hyperactivity leading to lower D-serine levels, and consequent hypo-excitation of NMDA receptors has been linked to schizophrenia risk. (43,44) We hypothesize that BCG-mediated *Dao*-repression (Fig. 5F, Supplementary File S1) may increase brain D–serine availability, boosting NMDA receptor co–agonist occupancy and improving synaptic plasticity and interneuron function in the PS19 mice.

Reduced p-tau levels in BCG-treated PS19 mice can be due to the repression of human tau (h-tau) P301S-expression. (31) To test this, we measured mRNA levels of h-tau P301S in the hippocampus tissue of PS19 females and the age-matched controls (Fig. 5A). We observed overexpression of h-tau P301S by ≈3-fold both in sham-treated and BCG-treated PS19 mice relative to age-matched non carrier controls. (Fig. 5G). Native murine tau (m-tau) levels were also unchanged in all the experimental groups (Fig. 5G), indicating similar expression of tau in the PS19 hippocampus. Additionally, bulk-RNA sequencing data suggested resolution of neuroinflammation in the BCG-treated PS19 mice. To test this, we quantified the expression of proinflammatory cytokines in hippocampal RNA from BCG-treated PS19 mice, and indeed we observed that the mRNA levels of *Ifnβ, Tnfα, Il1α, Il1β, Ccl8 and Mmp12* were significantly reduced relative to sham-treated PS19 mice and were restored to the levels of age-matched non carrier controls (Fig. 5H, 5I).

### BCG treatment reduces the activated microglial subpopulation without altering total microglial abundance in the PS19 brain

With the above data that there is no reduction in tau gene-expression, we turned to seeking mechanisms of increased clearance to explain the reduced p-tau depositions in the BCG-treated PS19 brains. Indeed, a comparison of the BCG-treated PS19 versus the AMC transcriptomes by GSEA found that the top cell type-specific DEGs were: ependymal cells (NES=2.96, *p.adj.=* 3.24E-30), dendritic cell (NES=2.81, *p.adj.=* 2.85E-15), macrophages (NES=2.68, *p.adj.=* 4.7E-16) and monocytes (NES=2.63, *p.adj.=* 7.59E-16). Moreover, “Antigen presentation and presentation of peptide antigen” was the top-enriched pathway (Fig. 5D, Supplementary file S1), suggesting that BCG induces a transcriptional program in myeloid cells consistent with increased capacity for uptake, lysosomal processing, and presentation of antigens- possibly tau-derived peptides- which may facilitate more efficient clearance of pathological tau aggregates.

To test this hypothesis, we isolated CD11b+ cells from whole mouse brains of 14-month-old sham-treated PS19, BCG-treated PS19 and age-matched control mice (Fig. 6A, fig. S5). Previous studies have reported that resting microglia under homeostatic conditions are CD11b^+^CD45^lo/int^. (45,46) However, under neuroinflammatory and neurodegenerative conditions, this tissue-resident microglial population acquires an activated CD11b+CD45^hi^ phenotype, making the distinction from infiltrating blood-derived/peripheral myeloid cells (CD11b+CD45^hi^) difficult. (45–47) Therefore, to separate these populations under heightened inflammatory conditions such as those likely to be present in PS19-tauopathy, we added the CD49d marker (Integrin alpha 4; *Itga4*) which is uniquely present in peripheral myeloid cells. (48,49) With these markers we were able to define the following brain cell populations: microglia: CD11b+ CD49d- Cd45+^(lo-hi)^, activated microglia: CD11b+CD49d-CD45^hi^, and infiltrating peripheral myeloid cells: CD11b+CD49d+CD45^hi^ (Fig. 6B, fig. S5). Gating of the CD11b+ parent population (and all other markers used) was established using a CD11b or respective fluorescence-minus-one (FMO) control to ensure specificity and avoid false-positive inclusion of non-myeloid cells. Samples from two mice were pooled per measurement to ensure sufficient cell yield with a total of n=6 mice per group. (Fig. 6A)

**Fig. 6.**
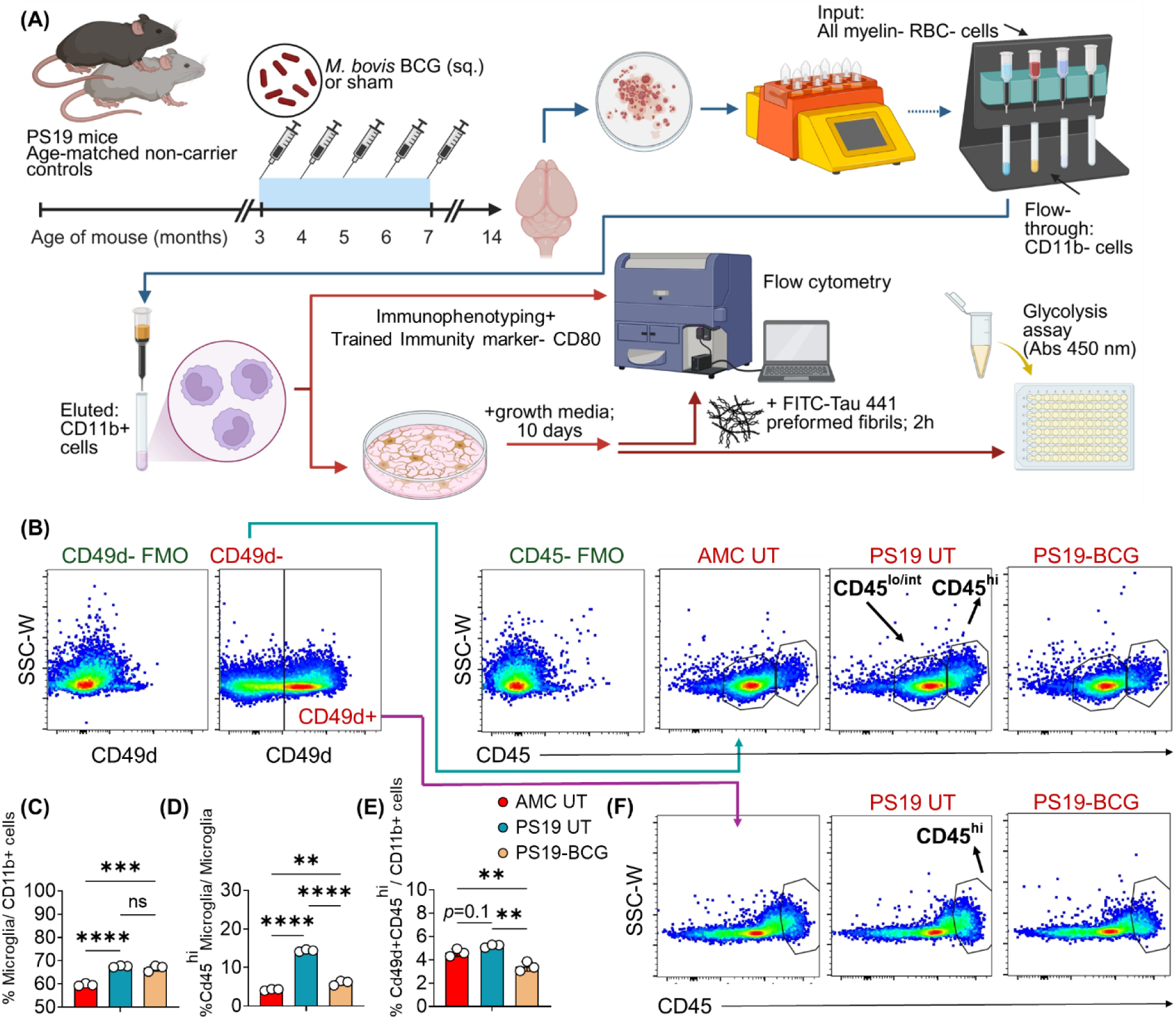
*M. bovis* BCG treatment reduces frequency of proinflammatory, activated microglial population, without reducing their total numbers in PS19 mice. (A) Schematic of the experiment. Magnetic bead-based enrichment of CD11b+ cells from all mouse brain cells (input). CD11b-FMO (fluorescence minus one) was used to set gating of CD11b+ population. (n=6 mice/group; samples from 2 mice were pooled together) **(B)** Microglia are defined as CD11b+CD49d-Cd45+ (low-high expression). **(C)** %microglial cells out of all CD11b+, **(D)** %activated microglia (CD45 high (hi) out of all microglia cells, and **(E)** %peripheral myeloid cells out of all CD11b+ cells in age-matched controls, sham-treated and BCG-treated PS19 mice, as determined by FlowJo software. **(F)** peripheral myeloid cells are CD11b+CD49d+Cd45high(hi). The data are means ± SEM, and representative of two independent experiments. Each data point represents a mouse. Statistical analysis was calculated by one-way ANOVA with Tukey’s multiple comparisons test. (*p*>0.05: ns, *p*<0.01: **, *p*<0.001: ***, *p*<0.0001: ****)

As expected, relative to age-matched controls, sham-treated PS19 mice have a significantly increased microglial population (*p*<0.0001) (Fig. 6C), and also a higher disease-associated CD45^hi^ microglia subpopulation (CD45^hi^ out of all CD45+ cells= 14.4% relative to 4.2% in AMC; *p*<0.0001) (Fig. 6D). In contrast, while we observed a similar increase in the total microglia population in the BCG-treated PS19 mice (Fig. 6C), there was significant reduction in the proportion of proinflammatory, activated microglia (CD45^hi^ out of all CD45+ cells= 6.04% relative to 14.4% in PS19 UT; *p*<0.0001), indicating normalization of the microglial activation state by BCG treatment (Fig. 6D). This flow cytometry data is agreement with the IHC data (Fig. 1C) where we observe a similar decrease in Iba1 levels in the BCG-treated PS19 hippocampus. In addition to this, the peripheral myeloid cells fraction increased in the sham-treated PS19 (5.1% out of all CD11b+ cells) relative to AMC (4.6%), but it was not statistically significant (*p*=0.1) (Fig. 6E, 6F). In contrast, the BCG-treated PS19 brains exhibited decreased infiltration of peripheral myeloid cells (3.4%), probably due to dampened neuroinflammation. Taken together, these data suggests that BCG therapy recalibrates the microglial composition in the brain—maintaining high microglial abundance while selectively dampening their proinflammatory state—thus reducing tissue–damaging inflammation (Fig. 5H, I) yet preserving a workforce to remove p-tau-deposits.

### BCG therapy induces trained-immunity features in brain myeloid cells

We hypothesized that AD prevention after BCG-therapy may share features with BCG-mediated myeloid cell “trained immunity”. (15–17) Previous studies have established that BCG-exposure “trains” central as well as peripheral immune cells by inducing both epigenetic (H3K4me3 and H3K27ac activation marks on immune-associated genes), metabolomic changes (increased glycolysis) and upregulation of pattern recognition receptors (*Tlr*), and costimulatory molecules (CD80, CD86). (50–52) The “training” occurs after the cell encounters a first stimulus (BCG in our study) and leads the cell to then display an elevated response against an unrelated, heterologous second stimulus (p-tau deposits in our study). To test this hypothesis, we measured the median fluorescence intensity of the CD80 surface marker on microglial and peripheral myeloid cells. Additionally, we quantified the phagocytosis and glycolysis levels of isolated CD11b+ cells (53,54). The CD80 levels on both microglial and peripheral myeloid cells from sham-treated PS19 mice were higher relative to age-matched controls (Fig. 7A, 7B), probably due to heightened inflammation. Interestingly, we observed higher expression of CD80 on both myeloid cell-types, but the peripheral myeloid cells from the BCG-treated PS19 mice exhibited increased expression of CD80 relative to both sham-treated PS19 and age-matched controls (*p*<0.05), suggesting compartment-specific induction of trained immunity. (Fig. 7A, 7B)

**Fig. 7.**
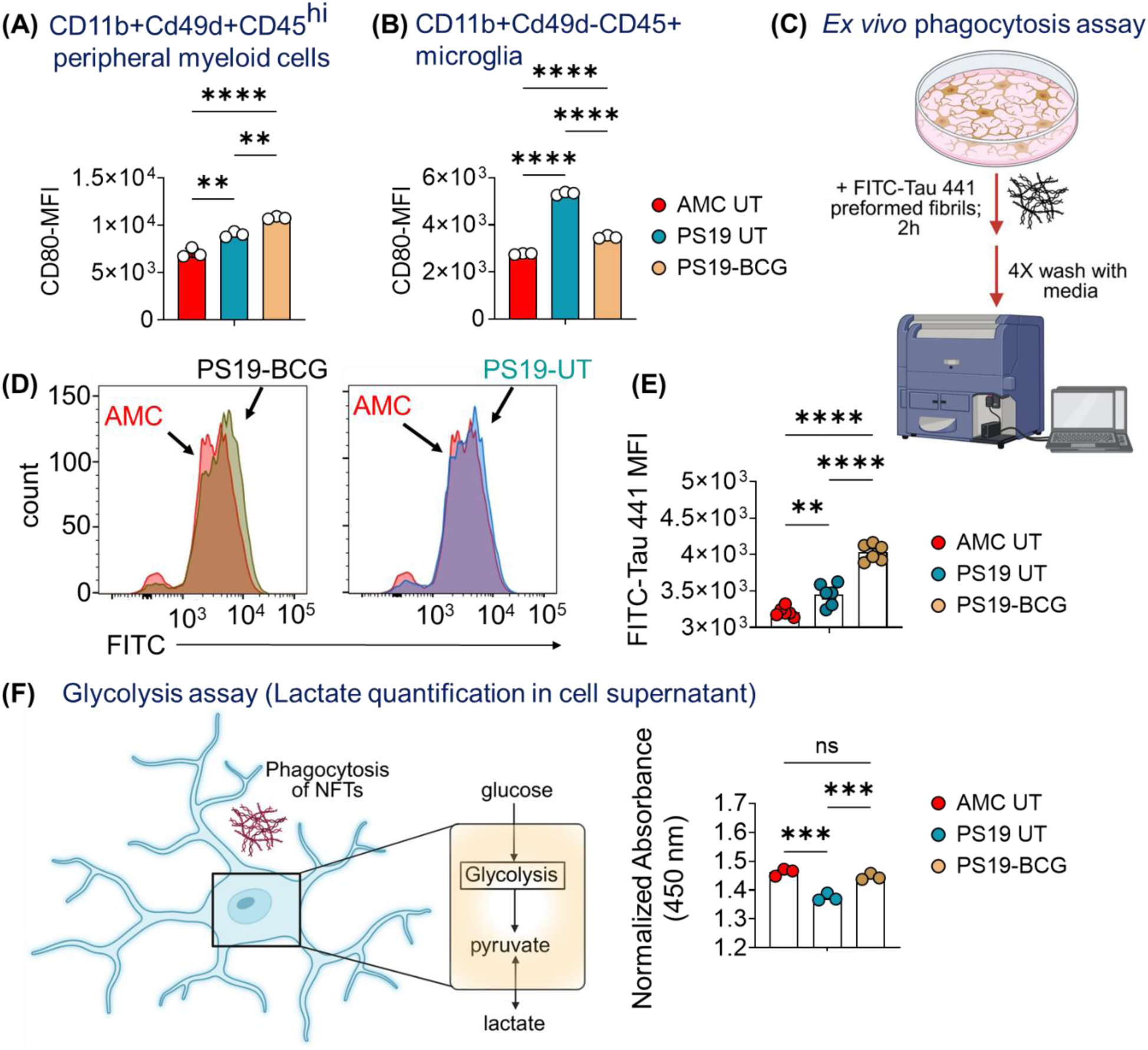
CD11b+ myeloid cells from BCG-treated PS19 mice brain displayed significantly higher markers of Trained Immunity. Median fluorescence intensity (MFI) of the surface marker of Trained immunity- CD80 on **(A)** peripheral myeloid cells and **(B)** microglia, as determined by FlowJo software. (n=6 mice/group; samples from 2 mice were pooled together) **(C)** Schematic of the phagocytosis assay with primary CD11b+ cells. **(D)** Histogram showing FITC-tau uptake by microglia, and respective **(E)** quantification. **(F)** Lactate concentration in the supernatant of resting CD11b+ cells, as determined by lactate dehydrogenase based colorimetric assay. The absorbance at 450 nm were normalized to respective protein levels. The data are means ± SEM, and representative of two independent experiments. Statistical analysis was calculated by one-way ANOVA with Tukey’s multiple comparisons test. (*p*>0.05: ns, *p*<0.01: **, *p*<0.001: ***, *p*<0.0001: ****)

Next, we measured the phagocytic capacity of primary brain CD11b+ cells *ex vivo,* using a physiologically relevant substrate, FITC-conjugated h-tau-441 fibrils. Adherent, primary brain CD11b+ myeloid cells were expanded in growth media for 10 days. They were then co-incubated with FITC-h-tau fibrils for 2 hours, extensively washed to remove extracellular fibrils, and then subjected to flow cytometry (Fig. 7C). While we observed a significantly high uptake of tau-fibrils by CD11b+ cells from sham-treated PS19 mice (7.5% higher relative to AMC; *p*=0.007), the CD11b+ cells from BCG-treated PS19 mice exhibited the highest uptake of the fibrils (25.4% higher relative to AMC; *p*<0.0001) (Fig. 7D, 7E) These data indicate that BCG-trained brain myeloid cells have an enhanced capacity for phagocytosis of p-tau-related protein deposits. In agreement with trained phenotype, CD11b+ brain myeloid cells from BCG-treated PS19 displayed higher production of lactate relative to sham-treated PS19 mice, (Fig. 7F) suggesting an elevated glycolysis setpoint at their baseline/resting state.

In sum, our results suggest that BCG therapy in the PS19 model shows features of myeloid cell immune training, that BCG therapy restricts microglial hyperactivation/microgliosis, while priming peripheral and/or brain myeloid cells for heightened p-tau–NFT phagocytosis, thereby mitigating tauopathy in the PS19 mice (Fig. 8).

**Fig. 8.**
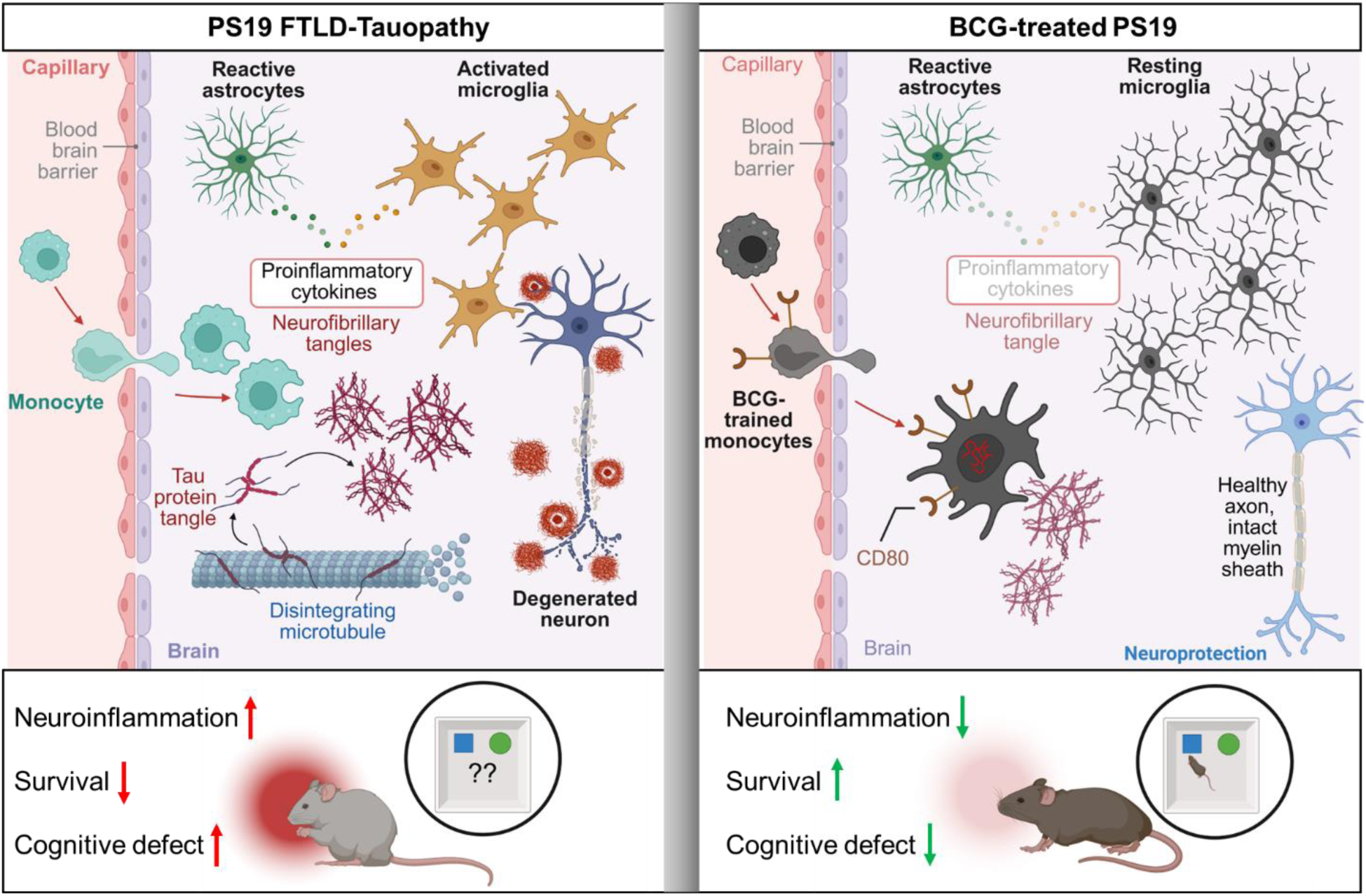
BCG-trained myeloid cells reduce neuroinflammation and neurodegeneration in PS19 tauopathy mice. Left: PS19 mice brains display blood–brain barrier dysfunction with infiltrating monocytes, reactive astrocytes, and activated microglia that release pro–inflammatory cytokines. Pathology includes tau aggregation into neurofibrillary tangles, and neuronal degeneration, accompanied by heightened systemic inflammation and cognitive/behavioral deficits. Right: BCG treatment induces trained monocytes (CD80+), which traffic to the brain, phagocytose and degrade NFTs, dampen microglial activation, and reduce pro–inflammatory cytokines. This is associated with fewer p-tau tangles, preserved axons and myelin, and overall neuroprotection, leading to reduced inflammation and improved cognitive performance in PS19 mice.

## Discussion

Our study is the first to assess BCG in a dementia model of tau pathology and to identify mechanisms by which BCG may promote clearance of pathologic tau NFTs. We found that early BCG administration starting at 3 months of age reduced brain atrophy, p-tau levels, and neuronal loss measured at the later time point of 8-9 months. Simultaneously, BCG therapy restored glutamate levels *in vivo*, improved recognition-memory performance, body weight, and survival of PS19 mice, suggesting key disease modifying effects.

One of the most remarkable effects of BCG that we observed in this study came from RNA sequencing of the hippocampus of 9.5 month-old mice revealing restoration of the transcriptomic landscape to near normal levels in BCG-treated PS19 mice (Fig. 5B); this effect was mediated by an intervention that was both physically remote (hypodermis to brain) and temporally remote (4.5 month time interval). Our RNA-sequencing analysis provided a valuable overview of BCG-mediated changes on the hippocampal transcriptional landscape; moreover, it served as a useful hypothesis generating tool, leading us to several testable postulates such as: (i) whether BCG-treatment of PS19 mice may stimulate neuronal survival by downregulating inhibitory synaptic gene expression, (ii) whether BCG may promote neuronal differentiation via downregulation of *Hes3*, or (iii) whether BCG may boost NMDA synaptic signaling via *Dao* downregulation and increased D-serine availability.

Even though large-scale prospective studies have yet to show definitively whether BCG does or does not prevent dementia, there has been literature speculation around potential mechanisms of BCG neuroprotection. These include (i) CNS anti-inflammatory activity,

(ii) immunometabolic reprogramming, (iii) epigenetic reprogramming, (iv) antimicrobial against heterologous pathogens such as HSV-1 or *P. gingivalis* that might drive or contribute to dementia risk. (55) The latter three mechanisms are each properties associated with BCG-mediated trained immunity, and can be readily demonstrated with *in vitro* systems, (56,57) animal models, (58) and human BCG intervention studies. (59)

Mechanistically, our study identified BCG-induced trained immunity properties as playing key roles in the clearance of tau NFTs. Using the surface co-stimulatory molecule CD80 as a marker of trained immunity, we found that BCG-vaccination led to a significant increase in the recruitment of CD80+ peripheral myeloid cells to the brain where they are likely to play a role in removal of NFT debris. In addition, we found that adherent CD11b+ myeloid cells from BCG-treated PS19 mice were significantly more active in phagocytosis of h-tau-441 fibrils and in the generation of lactate (glycolytic metabolomic shift), both known traits of trained immune myeloid cells. Lastly, we found that BCG mechanistically reduces the inflammatory state of brain microglial cells. While both treated and untreated PS19 mice had significantly elevated populations of brain microglial cells compared to healthy AMC mice, a much higher proportion of those microglial cells were activated in untreated PS19 mice (microgliosis) than in BCG-treated PS19 mice. The mechanisms of BCG action in the PS19 model are summarized diagrammatically in Fig. 8.

While our study is unique in addressing BCG in a tauopathy model of dementia, others have tested the efficacy of BCG for AD in the APP/PS1 β-amyloid (Aβ) mouse model and demonstrated improved cognition in BCG-treated APP/PS1 mice compared to untreated animals. (60–62) Across these three studies there were confounding results on the level of reduction (or lack thereof) of Aβ plaque formation, CNS anti-inflammatory cytokine levels (IL4, IL10, or TGFβ), and neurotrophic factors (BDNF). Recruitment of IL10-producing monocytes were shown to increase in the areas of Aβ plaques. (60–62) A concern with these studies is that brain atrophy correlates closely with p-tau accumulation, and less with Aβ deposition. (20,63–65) Our study also has certain limitations. The p-tau-driven neurodegeneration observed in the PS19 model most closely parallels the human dementia syndrome of frontotemporal lobar degeneration with tau pathology (FTLD-Tau) rather than classical human AD, which also includes β-amyloid deposition. The sex-differences observed in the PS19 model (females survive longer) are also a limitation; it is not feasible to compare sex- and age-matched animals since the disease is far more advanced in males at any given time point.

There is an urgent need for Alzheimer’s disease (AD)-modifying drugs since the three FDA-approved agents currently slow but not stop dementia progression, and neither cure nor proven prevention has been achieved. (66–69) AD belongs to the broader class of tauopathy syndromes which are characterized by the progressive accumulation and spread of misfolded, hyperphosphorylated tau fibrils leading to synaptic failure, neuronal death, and cognitive decline. (70) Histologically, AD also involves β-amyloid plaque deposition which precedes tau aggregation. (71,72) Given the failure of anti-amyloid therapies in clinical trials and the strong clinical correlation between NFT accumulation with AD severity in humans, (63,73–76) novel strategies ameliorating tauopathy may mitigate AD onset.

BCG is a live attenuated bacterial strain developed by serial passage of *Mycobacterium bovis* from 1908-1921, (77) and may be the most widely administered vaccine in human history, with an estimated 290 million doses administered to newborns annually to prevent tuberculosis. (78–81) BCG is also a critical immunotherapy for bladder cancer and is considered first line therapy against non-muscle invasive bladder cancer (NMIBC) which is the most common form of bladder cancer with an incidence ∼61,500 cases per year in the U.S. (17,82) BCG is also considered one of the world’s safest vaccines. Due to over one century of human use, adverse effects (AEs) of BCG are exceptionally well-defined. Local injection site reactions (pain, swelling) and flu-like symptoms occur at rates comparable to other vaccines. (83) Severe AEs—disseminated BCG infection (0.1-0.2 per 100,000) (84) and osteitis (0.3-1.0 per 100,000) (85)—are very rare.

While other vaccines such as the shingles vaccines, Zostavax and Shingrix, may also have dementia-preventative activity, these and most other vaccines that may have dementia-protective activity are already in use in the United States and most developed countries. Uniquely, BCG is a safe, readily-manufactured vaccine that is not used in the United States, Canada, Australia, New Zealand, and most western European nations (due to a low-incidence of TB)—the very same populations that bear the highest burden of reported-AD and related dementias. Our study strongly suggests that dementia protection by BCG is a credible observation that merits future basic research and large-scale prospective clinical trials to provide definitive answers.

## Materials and Methods

### Animals used in this study

The PS19 mouse line (B6;C3-Tg(Prnp-MAPT*P301S)PS19Vle/J) were purchased from The Jackson Laboratory (6 weeks old; Strain: #008169 (Bar Harbor, ME, USA)). This mouse strain was bred and maintained at the Rodent Vivarium of Johns Hopkins University, School of Medicine and was used in this study. The breeding strategy was to mate noncarrier × hemizygote, and genetic identity was confirmed by PCR-based genotyping. The oligonucleotides used for the PCR are shown in Supplementary file S2. All non-carrier age-matched controls used were littermates of hemizygous PS19 mice. All experiments were conducted under protocols approved by the Johns Hopkins Animal Care and Use Committee (Protocol Number: MO23M47) and followed established ethical animal handling and care guidelines. Animals were housed in individually ventilated cages under controlled conditions, including a 12-hour light/dark cycle, ambient temperatures ranging from 20 to 24°C, and relative humidity between 45% and 65%. Each cage contained no more than five mice of the same strain/sex to ensure proper animal welfare. Mice were randomly assigned to experimental groups to minimize bias. Humane and painless euthanasia procedures were employed in compliance with AVMA (American Veterinary Medical Association) guidelines.

All experiments involving *Mycobacterium bovis* BCG were conducted in Biosafety Level 2 (BSL2) facilities at Johns Hopkins University, School of Medicine, following protocols approved by the Institutional Biosafety Committee. Mice were briefly restrained during subcutaneous injection to ensure accurate delivery. All animals were monitored for signs of illness, such as weight loss or lethargy, and those that became moribund before the experiment’s conclusion were promptly euthanized, except in survival studies.

### Bacterial culture and subcutaneous administration into mice

*Mycobacterium bovis* BCG Pasteur TMC 1173P2 was obtained from the Johns Hopkins University, Center for Tuberculosis Research (CTBR). BCG was grown on 7H11 agar (Becton, Dickinson and Company, USA) supplemented with OADC (BD; 212351) at 37°C. CFU enumeration was performed by plating serial dilutions on 7H11 agar supplemented with OADC, followed by incubation at 37°C for 3 to 4 weeks.

For subcutaneous vaccination of mice, BCG cryostocks in Freezing mix (7H9+ 15% glycerol) (OD=1.2) were thawed, cells were washed twice with sterile 1X PBS and resuspended in 1X PBS. BCG strain was administered into mouse in a total volume of 0.2 ml per treatment. PS19 mice were either injected with phosphate buffered saline (1X PBS; sham) or BCG Pasteur strain (2x 10^6^ colony-forming units (CFU)) subcutaneously into the dorsal neck (scruff) region of mice. All mice were monitored daily for signs of tauopathy progression-including weight loss, paresis, lethargy, hunched posture, and reduced activity. Survival data were analyzed using Kaplan-Meier survival curves to determine the median time to death and assess differences between treatment groups.

### Immunohistochemistry against p-tau, Iba1, NeuN and GFAP

At indicated time points, the mice were euthanized by isoflurane (VetOne, Fluriso, MWI Animal Health, Boise, ID, USA) overdose. The mouse brains were harvested after transcardial perfusion with ≈15 mL of ice-cold 1X PBS (Gibco, Cat. No.-10010) and dissected on a chilled surface into two halves from midline. The left brain was dissected into hippocampus and rest of the brain, flash frozen in liquid nitrogen and stored at −80 °C, until tissue processing. The right half of the brain was fixed in 4% paraformaldehyde solution (Paraformaldehyde (Sigma-Aldrich, Cat. No.-P6148) in 1X PBS) for at least 24 hours at 4 °C, before it was submitted to the SKCCC Oncology Tissue and Imaging Service Core Laboratory (Johns Hopkins University, School of Medicine) for downstream processing. The brains were paraffin-embedded and 10 µm-sectioned.

For immunohistochemistry, the slides were deparaffinized and antigen retrieval was performed using 10mM sodium citrate buffer pH=6, to efficiently expose the epitope to the antibodies. Nonspecific binding of antibodies was eliminated by incubating with blocking buffer (1.5% normal goat serum (Vector Laboratories, Cat. No. S-1000-20) in PBS with 0.1% Triton X-100) for 1 hour. Slides were then incubated in 0.3% H_2_O_2_ (Sigma, Cat. No-H1009) for 30 minutes to quench endogenous peroxidase. Primary antibodies were prepared in blocking buffer and applied overnight at room-temperature (RT) followed by incubation with biotinylated secondary antibody (Vector Laboratories, Anti-mouse Ref. BP-9200-50; Anti-rabbit Ref. BP-9100-50) for 1 hour. Peroxidase labeled ABC reagent (Vector Laboratories, Newark, CA, USA) was applied for 30 minutes followed by signal development using 3,3 diaminobenzidine (DAB) (Vector Laboratories). Slides were counterstained with hematoxylin, dehydrated, cleared and mounted. The following primary antibodies were used: antiserum against NeuN (1:1,000; MAB377, Merck), phosphorylated tau AT8 (1:1,000; MN1020, Invitrogen), GFAP (1:1000; Z0334, Dako Cooperation), and Iba1 (1:1000; EPR16588 (ab178846), Abcam). Manufacturer-validated antibodies with demonstrated specificity were used. For each mouse brain sample, IHC was done in three planes (10 µm thick sections) for all the antibodies.

Slides were digitally scanned at 40× using a Hamamatsu Nanozoomer S210 digital slide scanner (Hamamatsu Photonics, Shizuoka, Japan) at 40X magnification (0.23 microns/pixel). Image files were submitted in https://digital.pathology.johnshopkins.edu/ using Concentriq LS platform (Proscia, PA, USA). %DAB-positive area was quantified as a percentage of the total region of interest, using an in-house protocol (fig. S2) developed with ImageJ software version 1.54m (NIH, USA) to minimize human bias, through standardized software-based settings.

### Behavioral tests

Behavioral assessments commenced following 5 months of vehicle or BCG vaccination treatment (n=8 mice/group). Female mice were transferred to a reversed light cycle room (12:12 h dark:light) 2 weeks prior to testing to allow circadian rhythm adaptation. All tests were conducted during the dark phase under dim red lighting. To minimize cumulative stress and carryover effects, only one test was performed per day, arranged in order of increasing stress intensity: Open Field Test (OFT), and Novel Object Recognition Test (NORT). Prior to each test, mice were acclimatized for 30 minutes in their home cages within the testing room. All apparatus were thoroughly cleaned with Vimoba disinfectant before each session and between animals to eliminate residual odor cues.

### Open field test (OFT)

The OFT was conducted to assess locomotor activity. Mice were individually placed in the center of a square open field arena (40 × 40 × 40 cm) and allowed to explore freely for 30 minutes. Locomotor activity was recorded using a San Diego Instruments Photobeam Activity System–Open Field (PAS-Open Field), equipped with a 16 × 16 photobeam frame with 1-inch beam spacing surrounding the enclosure. Parameters measured included total beam breaks as an index of general locomotor activity.

### Novel Object Recognition Test (NORT)

The NORT was conducted to assess recognition memory. The test was performed in an open field arena (40 × 40 × 40 cm) over two consecutive days, consisting of three phases: habituation, training, and testing. On the first day (habituation phase), mice were placed individually into the testing chamber without objects and allowed to explore freely for 10 minutes. After 24 hours, the training phase was conducted, in which two identical objects (matched for color, size, material, and shape) were placed along the diagonal of the chamber. Mice were allowed to explore both objects freely for 10 minutes. Following a 60-minute inter-trial interval, the test phase was conducted, in which one familiar object was replaced with a novel object that differed only in shape. Mice were allowed to explore freely for 5 minutes. The position of the novel object was counterbalanced across subjects to avoid location preference bias. Animal behavior was recorded using an overhead video camera, and object exploration duration was quantified by the blinded observers. The recognition index (RI) was calculated as the proportion of novel object exploration time relative to total exploration time. The discrimination index (DI) was calculated as: (Novel object exploration time − Familiar object exploration time) / Total exploration time.

### T2- weighted MRI

Acquisition: MRI experiments were conducted on a simultaneous 7T/30 PET/MRI scanner (Bruker BioSpin) equipped with a 72 mm inner-diameter transmit/receive volume coil and a 1H receive-only mouse brain surface coil. Mice were anesthetized with isoflurane and maintained under anesthesia throughout image acquisition with continuous respiratory monitoring. After localizer, anatomical brain MRI were acquired using axial 2D TurboRARE T2-weighted sequence with the following parameters: repetition time (TR) = 4000 ms, echo time (TE) = 20 ms, field of view (FOV) = 22 × 22 mm², matrix size = 256 × 256, which yielded in plane resolution 0.086X0.086 mm² resolution, slice thickness was 0.3 mm with 12 signal averages.

Analysis: After acquisition, DICOM images were analyzed using either ITK-SNAP 4.4.0 software (32) or by VivoQuant 2020 (Invicro). With VivoQuant 2020, semiautomatic segmentation method was used to quantify ventricular and hippocampal volumes with the 3D Brain Atlas Tool. Ventricular CSF was segmented using a predefined threshold, while the hippocampus was delineated using an automatic anatomic segmentation tool. All segmentations were manually refined with reference to a digital mouse brain atlas. Three-dimensional models were then generated, and total ventricular and hippocampal volumes were calculated and recorded. Investigators were blinded to study group allocation during image analysis.

### Glutamate-weighted CEST-MRI

CEST MRI experiments were performed using a continuous-wave saturation pulse (1 s long, 3 μT in amplitude) followed by a fast spin-echo MRI readout. A total of 61 saturation frequency offsets were acquired between ± 6 ppm (equally spaced). An additional acquisition at 50 ppm was used for normalization. In plane field of view was 22 × 22 mm^2^ with a matrix size of 64 × 64 (in plane resolution 0.34 × 0.34 mm^2^), and slice thickness was 1.5 mm. Data was processed by first correcting for B_0_ field inhomogeneity by voxel-wise fitting of the bulk water peak and assigning it as 0 ppm. The amplitude of the glutamate peak at 3 ppm was mapped on a voxel-wise basis using MTRasym analysis(86) and then taking the mean MTRasym signal between 3.5 ppm to 2.5 ppm. All data was analyzed in python using in-house written scripts.

### Bulk RNA-Sequencing of mouse hippocampus

RNA was extracted from hippocampus tissue samples using the RNeasy Mini kit (Qiagen, Cat. No.- 74104) and quantity and quality assessed by Bioanalyzer (Agilent Technologies, Palo Alto, CA, USA). Depletion of ribosomal RNA was performed using the Qiagen Fast Select rRNA HMR kit (Qiagen, Cat. No.-334387) according to the manufacturer’s instructions. A stranded mRNA library was prepared using the Illumina TruSeq Stranded Total RNA kit (Illumina, Cat. No.- 20020596) according to manufacturer’s instructions. Sequencing was performed using the Novaseq 6000 X platform sequencer and the TruSeq Cluster Kit (Illumina), resulting in approximately 50 million paired end reads per sample. The samples were sequenced using a 2x150 Pair-End (PE) configuration. Illumina’s bcl2fastq v2.20.0 was used to convert BCL files to FASTQ files. Raw sequencing quality was assessed using FastQC v0.11.7, and adapter trimming and quality filtering were performed with fastp v0.24.0. The processed reads were aligned to the mouse reference genome (mm39) using RSEM v1.3.3, which generated BAM files along with gene and transcript level expression estimates. Differential expression analysis and principal component analysis were conducted using DESeq2 v1.44.0. Gene set enrichment analysis was performed with fgsea v1.30.0, using gene sets obtained from the Molecular Signatures Database (MSigDB). Raw data were deposited in NCBI’s Gene Expression Omnibus (GEO, accession number- GSE326626). Reviewers can access this data using the following token number- ixqlgiouvhqvpgv.

### Quantitative real-time PCR from mouse hippocampus

At the indicated time point, mouse hippocampus tissue was lysed in TRIzol (Thermo Fisher Scientific; 15596026), and RNA was extracted using Qiagen RNeasy Mini Kit (74104) and RNase-free DNase Set (Qiagen; 79254), following the manufacturer’s protocol. RNA purity and concentration were quantified using Nanodrop One (Thermo Fisher Scientific). 1 µg of RNA was used for cDNA synthesis using the iScript cDNA Synthesis Kit (BioRad; 1708891) and quantitative RT-PCR was done with QuantStudio 3 (Applied Biosystems; Thermo Fisher Scientific) using gene-specific primers (Supplementary file S2) and iTaq Universal SYBR Green Supermix (BioRad; 1725121). Expression of genes was normalized with Ct value for *Gapdh* expression (internal housekeeping control).

### Isolation of CD11b+ cells from mouse brain

Adult brain dissociation kit (Miltenyi Biotec, Cat. No.-130-107-677) was used to dissociate brain into single-cell suspensions, following the manufacturer’s instructions. Briefly, the mice were euthanized via isoflurane overdose, their brains were harvested following complete transcardial perfusion with 15 mL of chilled 1X D-PBS (ThermoFisher scientific, Cat. No.- 14287072), washed in D-PBS, cut into small pieces using scalpel, and collected in 1950 µl Enzyme mix 1 + 30 µl Enzyme Mix 2 containing gentleMACS C-tubes. The cut-tissue was dissociated using the gentleMACS Octo Dissociator (program-37C_ABDK_01), and single-cell suspensions were obtained by filtering dissociated tissue through 100 µm MACS Smartstrainers (Miltenyi Biotec, Cat. No.-130-110-916). This was followed by removal of myelin debris and red blood cells, following manufacturer’s protocol. All calculations were based on the original mass of brain= 400-500 mg. Single cells were collected in PB buffer (D-PBS+ 0.5% fetal bovine serum (Millipore Sigma, Catalog no.-F4135)), passed through 70 µm MACS Smartstrainer (Miltenyi Biotec, Cat. No.-130-098-462) followed by centrifugation for 10 minutes at 4°C, 350 xg. The strainer was prerinsed with buffer to maximize recovery. The cell pellets were gently resuspended in 90 µL ice-cold PB buffer per 1 × 10⁷ total cells by gentle pipetting, as recommended by the manufacturer. CD11b magnetic MicroBeads (Miltenyi Biotec, 130-093-634) were added at 10 µL per 1 × 10⁷ cells, mixed by gentle inversion, and incubated for 15 min at 2–8°C in the dark. The cells were washed with 1 mL ice-cold PB buffer and centrifuged at 300 ×g for 5 min; supernatants were aspirated completely. Pellets were resuspended in 1000 µL PB buffer, and 20 µL of the suspension was reserved for subsequent flow cytometric analysis as the input (pre-enrichment) fraction. For magnetic separation, LS columns (Miltenyi Biotec, 130-042-401) with prerinsed 70 µm pre-separation filter were placed in a QuadroMACS Separator. Columns were equilibrated with 3 ml PB buffer prior to sample loading. The labeled cell suspension was applied to the column, and the flow-through was discarded. Columns were washed with PB buffer (3 times with 3 mL), removed from the magnetic field, placed over a clean collection tube, and magnetically retained cells were eluted with a plunger to obtain the CD11b+ fraction in PB buffer. Both fractions were centrifuged at 300 × g for 10 min at room temperature, supernatants were completely aspirated, and pellets were resuspended in 1 mL PB buffer. These cells were used immediately for downstream assays.

### Immunoprofiling of brain tissue by Multicolor Flow cytometry

Primary CD11b+ cells were counted by Cell counter (Invitrogen Countess Automated Cell Counter; C10281). We isolated ≈2x 10^5^ CD11b+ cells/ mouse brain and samples from 2 mouse samples were pooled together. The cells were centrifuged at 300 ×g for 5 minutes, supernatant was aspirated out completely and resuspended in 1X PBS. Primary cells were stained for viability using a live-dead near-IR dye (1:1000 dilution in PBS; Thermo Fisher Scientific; L10119) and blocked with Rat Anti-Mouse CD16/CD32 Fc Block (Clone 2.4G2, BD Pharmingen; 553142). The panel of antibodies was designed using FluoroFinder (https://fluorofinder.com/). For extracellular staining, cells were incubated with 50 µL of an antibody cocktail in Brilliant Stain Buffer (BD Biosciences; 563794) containing anti-mouse CD45 (Brilliant Violet 480; BD Biosciences;566095), CD11b (Brilliant Violet 650; BioLegend; 101259), CD49d (APC; BioLegend; 103622), CD80 (B7-1) monoclonal antibody (16-10A1; eFluor 450; eBioscience; 48-0801-82) for 30 minutes at 4°C in the dark. After staining, cells were fixed with 4% paraformaldehyde solution, washed with Cell staining buffer (BioLegend; 420201), filtered through strainer-capped BD FACS tubes (BD; 352235), and analyzed via flow cytometry. 1×10^5^ events per sample were acquired using a BD LSR Fortessa™ Cell Analyzer (BD Biosciences), and the data were analyzed with FlowJo software version 10.8.1 (BD Biosciences). Single stain controls, Fluorescence minus one (FMOs) and compensation matrices were prepared using cells. All flow antibodies were titrated to determine optimal staining concentrations, and only manufacturer-validated antibodies with demonstrated specificity were used. For the analysis, cells were first gated on singlets (FSC-H vs. FSC-A) and live cells (live-dead near-IR negative) before further selection of cell types. Microglia and peripheral myeloid cells were defined as: Cd11b+Cd45+CD49d-, and Cd11b+Cd45hiCD49d+ cells, respectively. Gating for all the positively-stained events were established based on respective Fluorescence minus one (FMO) and unstained controls.

### Primary cells culture and Phagocytosis assay

Primary CD11b+ cells were isolated from brain tissue as described above. Single cells were seeded at a density of 5×10^4^ cells/ well onto poly-D-lysine (PDL) (Sigma Aldrich; A003E)-coated 24-well plates in Dulbecco’s modified Eagle’s medium (DMEM, high glucose, GlutaMAX™; Thermo Fisher Scientific; 10566016) containing 10% heat-inactivated FBS (Millipore Sigma; F4135), 1X antibiotic-antimycotic Solution (Sigma Aldrich; A5955) and 100 ng/ml macrophage colony-stimulating factor (M-CSF; PeproTech, Cat.# 315-02). The cells were maintained at 37 °C in a humidified incubator with 5% CO_2_ for 10 days. (87) The following day, medium was replaced and cells were treated with 5 µg of FITC-conjugated h-tau-441 preformed fibrils (rPeptide; TF-1113-2) for 2 hours at 37°C in a humidified atmosphere with 5% CO_2_. After treatment, cells were washed twice with phosphate-buffered saline (PBS) and scraped out in 1ml RPMI (without phenol red) (Gibco; 11835). Live-dead Fixable Near-IR Dead Cell Stain (Thermo Fisher Scientific; L10119) was added for the detection of dead cells. FACS was immediately done using CytoFLEX cytometer (Beckman Coulter), and 10000 events/sample were recorded. The FlowJo software was applied for the quantification of cell population and fluorescence intensity.

### Glycolysis assay

The glycolysis capacity of CD11b+ cells were measured using the Glycolysis/OXPHOS Assay Kit (Dojindo, Tabaru, Japan; G270) according to the manufacturer’s instructions. Single cells were seeded at a density of 10^4^ cells/ well onto poly-D-lysine (PDL)-coated 96-well plates and maintained at 37 °C in a humidified incubator with 5% CO_2_ for 10 days in complete media. The following day, medium was replaced with fresh media and incubated further for 2 days. The lactate in the culture supernatant was quantified by incubating 20 µL of the culture medium with 80 µL lactate working solution at 37°C for 60 minutes, after which the absorbance at 450 nm was measured using the Bio-Rad iMark microplate reader. The absorbance at 450 nm was normalized to the respective protein concentration of sample (as measured by the BCA Assay kit; Thermo Scientific; 79023227).

### Statistical analysis

The sample size was not predetermined by statistical methods, but it is consistent with our previous publications. Statistical analyses were performed using GraphPad Prism version 10.4.0 software. First, Normality distribution was assessed using the Shapiro-Wilk test. Statistical significance was determined using unpaired two-tailed Student’s t-tests or Mann Whitney test for two-group comparisons, and one-way or two- way ANOVA with post hoc tests for multiple-group comparisons. All data points represent biological replicates; samples were not measured repeatedly to obtain data. Kaplan-Meier survival analysis was conducted for time-to-death (TTD) studies, with group differences assessed by the Log-rank (Mantel-Cox) test. In animal experiments, each data point represented a single mouse, and experiments were randomized, though blinding was not implemented. Mice that became moribund prior to the end of the experimental time period were sacrificed immediately, except for in survival studies. Results are presented as mean with error bars indicating the Standard Error of the Mean (SEM). *p*-values less than 0.05 were considered statistically significant.

### Contact for reagent and resource sharing

All data associated with this study are present in the paper- either in the Results section or in the Supplementary files. Requests for further details on dataset, resources, protocol, and reagents can be directed to the corresponding author- Prof. William R Bishai. Raw data generated during this study is provided with this paper as a separate Excel file labeled “Source Data”. The RNA sequencing dataset is available on NCBI’s Gene Expression Omnibus (GEO accession number- GSE326626).

## Acknowledgements

We gratefully acknowledge the support of the NIH grants: R01 AI155346, P30 CA006973, R01 CA281075, R01-AI190038. The funders have no role in study-design, data collection and interpretation, and the decision to submit the work for publication or preparation of the manuscript. We sincerely thank all the members of Prof. Bishai lab for their valuable suggestions and discussions throughout this project. We extend our gratitude to the JHU-SKCCC Core centers for their assistance with Histology and Flow cytometry, and MRB Molecular Imaging Service Center and Cancer Functional Imaging Core for MRI data acquisition. The schematic Figures in this study were created using BioRender (with Academic Lab License).

## Author contributions

Funding Acquisition was contributed by- WB, BS, PW. Conceptualization- WB, SS. Methodology- SS, MH, YZ, MSB, CRG, SY, SL, ST, SJ, NY, EN, ZB. Investigation and Formal Analysis- WB, SS, MH, MSB, SY, NY. Data curation and Validation- SS, MH, NY. Writing original draft- WB, SS. Review and editing manuscript- WB, PW, BS, SS, MH, YZ, MSB, CRG, SY, SL, ST, SJ, NY, EN, ZB.

## References

1. Eyting M, Xie M, Michalik F, Heß S, Chung S, Geldsetzer P. A natural experiment on the effect of herpes zoster vaccination on dementia. Nature 2025 641:8062. 2025 Apr 2;641(8062):438–46. doi:10.1038/s41586-025-08800-x PubMed PMID: 40175543.

2. Taquet M, Dercon Q, Todd JA, Harrison PJ. The recombinant shingles vaccine is associated with lower risk of dementia. Nature Medicine 2024 30:10. 2024 Jul 25;30(10):2777–81. doi:10.1038/s41591-024-03201-5 PubMed PMID: 39053634.

3. Rayens E, Sy LS, Qian L, Ackerson BK, Tubert J, Luo Y, et al. Recombinant zoster vaccine is associated with a reduced risk of dementia. Nature Communications 2026. 2026 Feb 9. doi:10.1038/s41467-026-69289-0

4. Xie M, Eyting M, Bommer C, Ahmed H, Geldsetzer P. The effect of shingles vaccination at different stages of the dementia disease course. Cell. 2025 Dec 11;188(25):7049–7064.e20. doi:10.1016/j.cell.2025.11.007 PubMed PMID: 41338191.

5. Gofrit ON, Klein BY, Cohen IR, Ben-Hur T, Greenblatt CL, Bercovier H. Bacillus Calmette-Guérin (BCG) therapy lowers the incidence of Alzheimer’s disease in bladder cancer patients. PLoS One. 2019 Nov 1;14(11):e0224433. doi:10.1371/journal.pone.0224433 PubMed PMID: 31697701.

6. Weinberg MS, Zafar A, Magdamo C, Chung SY, Chou WH, Nayan M, et al. Association of BCG Vaccine Treatment With Death and Dementia in Patients With Non–Muscle-Invasive Bladder Cancer. JAMA Netw Open. 2023 May 1;6(5):e2314336–e2314336. doi:10.1001/jamanetworkopen.2023.14336 PubMed PMID: 37204792.

7. Kim JI, Zhu D, Barry E, Kovac E, Aboumohamed A, Agalliu I, et al. Intravesical Bacillus Calmette-Guérin Treatment Is Inversely Associated With the Risk of Developing Alzheimer Disease or Other Dementia Among Patients With Non–muscle-invasive Bladder Cancer. Clin Genitourin Cancer. 2021 Dec 1;19(6):e409–16. doi:10.1016/j.clgc.2021.05.001 PubMed PMID: 34116955.

8. Klinger D, Hill BL, Barda N, Halperin E, Gofrit ON, Greenblatt CL, et al. Bladder Cancer Immunotherapy by BCG Is Associated with a Significantly Reduced Risk of Alzheimer’s Disease and Parkinson’s Disease. Vaccines 2021, Vol 9,. 2021 May 10;9(5). doi:10.3390/vaccines9050491

9. Makrakis D, Holt SK, Bernick C, Grivas P, Gore JL, Wright JL. Intravesical BCG and Incidence of Alzheimer Disease in Patients with Bladder Cancer: Results from an Administrative Dataset. Alzheimer Dis Assoc Disord. 2022 Oct 1;36(4):307–11. doi:10.1097/WAD.0000000000000530 PubMed PMID: 36183417.

10. Wang E, Hagberg O, Malmström PU. The association between BCG treatment in patients with bladder cancer and subsequent risk of developing Alzheimer and other dementia.-A Swedish nationwide cohort study from 1997 to 2019. PLoS One. 2023 Dec 1;18(12). doi:10.1371/journal.pone.0292174 PubMed PMID: 38096211.

11. Umar TP, Jain N, Stevanny B, Javed B, Priandhana A, Siburian R, et al. Protective role of Bacillus Calmette–Guérin vaccine in Alzheimer’s disease progression: A systematic review and meta-analysis. Heliyon. 2024 Mar 15;10(5):e27425. doi:10.1016/J.HELIYON.2024.E27425

12. Ibrahim M, Kim P, Marawar R, Avgerinos KI. Bacillus Calmette-Guerin (BCG) Vaccine Impact on Dementia Risk in Bladder Cancer Patients: A Systematic Review and Meta-Analysis. J Prev Alzheimers Dis. 2024 Oct 1;11(5):1355–62. doi:10.14283/jpad.2024.94 PubMed PMID: 39350381.

13. Han C, Wang J, Chen YL, Guan CP, Zhang YA, Wang MS. The role of Bacillus Calmette-Guérin administration on the risk of dementia in bladder cancer patients: a systematic review and meta-analysis. Front Aging Neurosci. 2023 Aug 24;15:1243588. doi:10.3389/fnagi.2023.1243588

14. Dow CT, Greenblatt CL, Chan ED, Dow JF. Evaluation of BCG Vaccination and Plasma Amyloid: A Prospective, Pilot Study with Implications for Alzheimer’s Disease. Microorganisms 2022, Vol 10,. 2022 Feb 11;10(2). doi:10.3390/microorganisms10020424

15. Netea MG, Joosten LAB, Latz E, Mills KHG, Natoli G, Stunnenberg HG, et al. Trained immunity: A program of innate immune memory in health and disease. Science. 2016 Apr 22;352(6284):aaf1098. doi:10.1126/science.aaf1098 PubMed PMID: 27102489.

16. van Puffelen JH, Keating ST, Oosterwijk E, van der Heijden AG, Netea MG, Joosten LAB, et al. Trained immunity as a molecular mechanism for BCG immunotherapy in bladder cancer. Nat Rev Urol. 2020 Sep 16;17(9):513–25. doi:10.1038/s41585-020-0346-4

17. Shee S, Bishai W. Induction of Trained Immunity and Nonspecific Protective Effects Against Heterologous Diseases by BCG Vaccination. Tuberculosis Vaccines. 2025 Jan 1;65–94. doi:10.1007/978-3-031-94540-3_4

18. Tiwari V, Prajapati B, Asare Y, Damkou A, Ji H, Liu L, et al. Innate immune training restores pro-reparative myeloid functions to promote remyelination in the aged central nervous system. Immunity. 2024 Sep 10;57(9):2173–2190.e8. doi:10.1016/j.immuni.2024.07.001 PubMed PMID: 39053462.

19. Yoshiyama Y, Higuchi M, Zhang B, Huang SM, Iwata N, Saido TCC, et al. Synapse Loss and Microglial Activation Precede Tangles in a P301S Tauopathy Mouse Model. Neuron. 2007 Feb 1;53(3):337–51. doi:10.1016/j.neuron.2007.01.010 PubMed PMID: 17270732.

20. Takeuchi H, Iba M, Inoue H, Higuchi M, Takao K, Tsukita K, et al. P301S Mutant Human Tau Transgenic Mice Manifest Early Symptoms of Human Tauopathies with Dementia and Altered Sensorimotor Gating. PLoS One. 2011;6(6):e21050. doi:10.1371/journal.pone.0021050 PubMed PMID: 21698260.

21. Cao Q, Kumar M, Frazier A, Williams JB, Zhao S, Yan Z. Longitudinal characterization of behavioral, morphological and transcriptomic changes in a tauopathy mouse model. Aging. 2023 Nov 3;15(21):11697–719. doi:10.18632/aging.205057 PubMed PMID: 37925173.

22. Leyns CEG, Ulrich JD, Finn MB, Stewart FR, Koscal LJ, Serrano JR, et al. TREM2 deficiency attenuates neuroinflammation and protects against neurodegeneration in a mouse model of tauopathy. Proceedings of the National Academy of Sciences. 2017 Oct 24;114(43):11524–9. doi:10.1073/pnas.1710311114 PubMed PMID: 29073081.

23. Shi Y, Manis M, Long J, Wang K, Sullivan PM, Serrano JR, et al. Microglia drive APOE-dependent neurodegeneration in a tauopathy mouse model. Journal of Experimental Medicine. 2019 Nov 4;216(11):2546–61. doi:10.1084/jem.20190980 PubMed PMID: 31601677.

24. Ising C, Venegas C, Zhang S, Scheiblich H, Schmidt S V., Vieira-Saecker A, et al. NLRP3 inflammasome activation drives tau pathology. Nature 2019 575:7784. 2019 Nov 20;575(7784):669–73. doi:10.1038/s41586-019-1769-z PubMed PMID: 31748742.

25. Maphis N, Xu G, Kokiko-Cochran ON, Jiang S, Cardona A, Ransohoff RM, et al. Reactive microglia drive tau pathology and contribute to the spreading of pathological tau in the brain. Brain. 2015 Jun 1;138(6):1738–55. doi:10.1093/brain/awv081 PubMed PMID: 25833819.

26. Gao C, Jiang J, Tan Y, Chen S. Microglia in neurodegenerative diseases: mechanism and potential therapeutic targets. Signal Transduction and Targeted Therapy 2023 8:1. 2023 Sep 22;8(1):359-. doi:10.1038/s41392-023-01588-0 PubMed PMID: 37735487.

27. Abellanas MA, Purnapatre M, Burgaletto C, Schwartz M. Monocyte-derived macrophages act as reinforcements when microglia fall short in Alzheimer’s disease. Nature Neuroscience 2025 28:3. 2025 Jan 6;28(3):436–45. doi:10.1038/s41593-024-01847-5 PubMed PMID: 39762659.

28. Sun Y, Guo Y, Feng X, Jia M, Ai N, Dong Y, et al. The behavioural and neuropathologic sexual dimorphism and absence of MIP-3α in tau P301S mouse model of Alzheimer’s disease. Journal of Neuroinflammation 2020 17:1. 2020 Feb 24;17(1):72-. doi:10.1186/s12974-020-01749-w PubMed PMID: 32093751.

29. Huang M, Tallon C, Zhu X, Huizar KDJ, Picciolini S, Thomas AG, et al. Microglial-Targeted nSMase2 Inhibitor Fails to Reduce Tau Propagation in PS19 Mice. Pharmaceutics 2023, Vol 15,. 2023 Sep 20;15(9). doi:10.3390/pharmaceutics15092364

30. Wu T, Dejanovic B, Gandham VD, Gogineni A, Edmonds R, Schauer S, et al. Complement C3 Is Activated in Human AD Brain and Is Required for Neurodegeneration in Mouse Models of Amyloidosis and Tauopathy. Cell Rep. 2019 Aug 20;28(8):2111–2123.e6. doi:10.1016/j.celrep.2019.07.060 PubMed PMID: 31433986.

31. DeVos SL, Miller RL, Schoch KM, Holmes BB, Kebodeaux CS, Wegener AJ, et al. Tau reduction prevents neuronal loss and reverses pathological tau deposition and seeding in mice with tauopathy. Sci Transl Med. 2017 Jan 25;9(374). doi:10.1126/scitranslmed.aag0481 PubMed PMID: 28123067.

32. Yushkevich PA, Piven J, Hazlett HC, Smith RG, Ho S, Gee JC, et al. User-guided 3D active contour segmentation of anatomical structures: Significantly improved efficiency and reliability. Neuroimage. 2006 Jul 1;31(3):1116–28. doi:10.1016/j.neuroimage.2006.01.015 PubMed PMID: 16545965.

33. Crescenzi R, DeBrosse C, Nanga RPR, Reddy S, Haris M, Hariharan H, et al. In vivo measurement of glutamate loss is associated with synapse loss in a mouse model of tauopathy. Neuroimage. 2014 Nov 1;101:185–92. doi:10.1016/j.neuroimage.2014.06.067 PubMed PMID: 25003815.

34. Crescenzi R, DeBrosse C, Nanga RPR, Byrne MD, Krishnamoorthy G, D’Aquilla K, et al. Longitudinal imaging reveals subhippocampal dynamics in glutamate levels associated with histopathologic events in a mouse model of tauopathy and healthy mice. Hippocampus. 2017 Mar 1;27(3):285–302. doi:10.1002/hipo.22693 PubMed PMID: 27997993.

35. Zheng Y, Huang M, Maragakis RM, Pietri P, Su Y, Alt J, et al. Targeting NAAG metabolism restores cognition and synaptic integrity in EcoHIV-infected mice. Neurotherapeutics. 2025 Nov 6;95(19):e00782. doi:10.1016/j.neurot.2025.e00782

36. Saul D, Kosinsky RL, Atkinson EJ, Doolittle ML, Zhang X, LeBrasseur NK, et al. A new gene set identifies senescent cells and predicts senescence-associated pathways across tissues. Nat Commun. 2022;13(1). doi:10.1038/s41467-022-32552-1 PubMed PMID: 35974106.

37. Subramanian A, Tamayo P, Mootha VK, Mukherjee S, Ebert BL, Gillette MA, et al. Gene set enrichment analysis: A knowledge-based approach for interpreting genome-wide expression profiles. Proc Natl Acad Sci U S A. 2005 Oct 25;102(43):15545–50. doi:10.1073/pnas.0506580102 PubMed PMID: 16199517.

38. Mootha VK, Lindgren CM, Eriksson KF, Subramanian A, Sihag S, Lehar J, et al. PGC-1α-responsive genes involved in oxidative phosphorylation are coordinately downregulated in human diabetes. Nature Genetics 2003 34:3. 2003 Jun 15;34(3):267–73. doi:10.1038/ng1180 PubMed PMID: 12808457.

39. Hatakeyama J, Bessho Y, Katoh K, Ookawara S, Fujioka M, Guillemot F, et al. Hes genes regulate size, shape and histogenesis of the nervous system by control of the timing of neural stem cell differentiation. Development. 2004 Nov 15;131(22):5539–50. doi:10.1242/dev.01436 PubMed PMID: 15496443.

40. Molla G, Sacchi S, Bernasconi M, Pilone MS, Fukui K, Pollegioni L. Characterization of human d-amino acid oxidase. FEBS Lett. 2006 Apr 17;580(9):2358–64. doi:10.1016/j.febslet.2006.03.045 PubMed PMID: 16616139.

41. Wolosker H, Blackshaw S, Snyder SH. Serine racemase: A glial enzyme synthesizing D-serine to regulate glutamate-N-methyl-D-aspartate neurotransmission. Proc Natl Acad Sci U S A. 1999 Nov 9;96(23):13409–14. doi:10.1073/pnas.96.23.13409 PubMed PMID: 10557334.

42. Verrall L, Burnet PWJ, Betts JF, Harrison PJ. The neurobiology of D-amino acid oxidase and its involvement in schizophrenia. Molecular Psychiatry 2010 15:2. 2009 Sep 29;15(2):122–37. doi:10.1038/mp.2009.99 PubMed PMID: 19786963.

43. Mothet JP, Parent AT, Wolosker H, Brady RO, Linden DJ, Ferris CD, et al. D-serine is an endogenous ligand for the glycine site of the N-methyl-D-aspartate receptor. Proc Natl Acad Sci U S A. 2000 Apr 25;97(9):4926–31. doi:10.1073/pnas.97.9.4926 PubMed PMID: 10781100.

44. Tsai G, Yang P, Chung LC, Lange N, Coyle JT. D-serine added to antipsychotics for the treatment of schizophrenia. Biol Psychiatry. 1998 Dec 1;44(11):1081–9. doi:10.1016/S0006-3223(98)00279-0 PubMed PMID: 9836012.

45. Honarpisheh P, Lee J, Banerjee A, Blasco-Conesa MP, Honarpisheh P, d’Aigle J, et al. Potential caveats of putative microglia-specific markers for assessment of age-related cerebrovascular neuroinflammation. Journal of Neuroinflammation 2020 17:1. 2020 Dec 1;17(1):366-. doi:10.1186/s12974-020-02019-5 PubMed PMID: 33261619.

46. Sedgwick JD, Schwender S, Imrich H, Dörries R, Butcher GW, Meulen V Ter. Isolation and direct characterization of resident microglial cells from the normal and inflamed central nervous system. Proc Natl Acad Sci U S A. 1991 Aug 15;88(16):7438–42. doi:10.1073/pnas.88.16.7438 PubMed PMID: 1651506.

47. Müller A, Brandenburg S, Turkowski K, Müller S, Vajkoczy P. Resident microglia, and not peripheral macrophages, are the main source of brain tumor mononuclear cells. Int J Cancer. 2015 Jul 15;137(2):278–88. doi:10.1002/ijc.29379 PubMed PMID: 25477239.

48. Manocha G, Ghatak A, Puig K, Combs C. Anti-α4β1 Integrin Antibodies Attenuated Brain Inflammatory Changes in a Mouse Model of Alzheimer’s Disease. Curr Alzheimer Res. 2018 Aug 1;15(12):1123. doi:10.2174/1567205015666180801111033 PubMed PMID: 30068274.

49. Bowman RL, Klemm F, Akkari L, Pyonteck SM, Sevenich L, Quail DF, et al. Macrophage Ontogeny Underlies Differences in Tumor-Specific Education in Brain Malignancies. Cell Rep. 2016 Nov 22;17(9):2445–59. doi:10.1016/j.celrep.2016.10.052 PubMed PMID: 27840052.

50. Jeljeli M, Riccio LGC, Doridot L, Chêne C, Nicco C, Chouzenoux S, et al. Trained immunity modulates inflammation-induced fibrosis. Nature Communications 2019 10:1. 2019 Dec 11;10(1):5670-. doi:10.1038/s41467-019-13636-x PubMed PMID: 31827093.

51. Murphy DM, Cox DJ, Connolly SA, Breen EP, Brugman AAI, Phelan JJ, et al. Trained immunity is induced in humans after immunization with an adenoviral vector COVID-19 vaccine. J Clin Invest. 2023 Jan 17;133(2). doi:10.1172/JCI162581 PubMed PMID: 36282571.

52. Takeuchi O, Akira S. In vitro protocol demonstrating five functional steps of trained immunity in mice: Implications on biomarker discovery and translational research. Cell Rep. 2025 Oct 28;44(10):116202. doi:10.1016/j.cell.2010.01.022 PubMed PMID: 20303872.

53. Cirovic B, de Bree LCJ, Groh L, Blok BA, Chan J, van der Velden WJFM, et al. BCG Vaccination in Humans Elicits Trained Immunity via the Hematopoietic Progenitor Compartment. Cell Host Microbe. 2020 Aug 12;28(2):322–334.e5. doi:10.1016/j.chom.2020.05.014 PubMed PMID: 32544459.

54. Jeyanathan M, Vaseghi-Shanjani M, Afkhami S, Grondin JA, Kang A, D’Agostino MR, et al. Parenteral BCG vaccine induces lung-resident memory macrophages and trained immunity via the gut–lung axis. Nature Immunology 2022 23:12. 2022 Dec 1;23(12):1687–702. doi:10.1038/s41590-022-01354-4 PubMed PMID: 36456739.

55. Devine J, Jacobs B, Leroux-Roels I, Leroux-Roels G, van der Most R. Infection, vaccination and risk of dementia: a proposed immunological model. Front Immunol. 2026 Mar 4;17:1748535. doi:10.3389/FIMMU.2026.1748535/TEXT

56. Saeed S, Quintin J, Kerstens HHD, Rao NA, Aghajanirefah A, Matarese F, et al. Epigenetic programming of monocyte-to-macrophage differentiation and trained innate immunity. Science (1979). 2014 Sep 26;345(6204). doi:10.1126/SCIENCE.1251086;PAGE:STRING:ARTICLE/CHAPTER PubMed PMID: 25258085.

57. Arts RJW, Novakovic B, ter Horst R, Carvalho A, Bekkering S, Lachmandas E, et al. Glutaminolysis and Fumarate Accumulation Integrate Immunometabolic and Epigenetic Programs in Trained Immunity. Cell Metab. 2016 Dec 13;24(6):807–19. doi:10.1016/J.CMET.2016.10.008 PubMed PMID: 27866838.

58. Kaufmann E, Khan N, Tran KA, Ulndreaj A, Pernet E, Fontes G, et al. BCG vaccination provides protection against IAV but not SARS-CoV-2. Cell Rep. 2022 Mar;38(10):110502. doi:10.1016/j.celrep.2022.110502

59. Arts RJW, Moorlag SJCFM, Novakovic B, Li Y, Wang SY, Oosting M, et al. BCG Vaccination Protects against Experimental Viral Infection in Humans through the Induction of Cytokines Associated with Trained Immunity. Cell Host Microbe. 2018 Jan 10;23(1):89–100.e5. doi:10.1016/j.chom.2017.12.010 PubMed PMID: 29324233.

60. Zuo Z, Qi F, Xing Z, Yuan L, Yang Y, He Z, et al. Bacille Calmette-Guérin attenuates vascular amyloid pathology and maximizes synaptic preservation in APP/PS1 mice following active amyloid-β immunotherapy. Neurobiol Aging. 2021 May 1;101:94–108. doi:10.1016/j.neurobiolaging.2021.01.001

61. Zuo Z, Qi F, Yang J, Wang X, Wu Y, Wen Y, et al. Immunization with Bacillus Calmette-Guérin (BCG) alleviates neuroinflammation and cognitive deficits in APP/PS1 mice via the recruitment of inflammation-resolving monocytes to the brain. Neurobiol Dis. 2017 May 1;101:27–39. doi:10.1016/j.nbd.2017.02.001

62. Li Q, Wang X, Wang ZH, Lin Z, Yang J, Chen J, et al. Changes in dendritic complexity and spine morphology following BCG immunization in APP/PS1 mice. Hum Vaccin Immunother. 2022 Nov 30;18(6):2121568. doi:10.1080/21645515.2022.2121568 PubMed PMID: 36113067.

63. Hanseeuw BJ, Betensky RA, Jacobs HIL, Schultz AP, Sepulcre J, Becker JA, et al. Association of Amyloid and Tau With Cognition in Preclinical Alzheimer Disease: A Longitudinal Study. JAMA Neurol. 2019 Aug 1;76(8):915–24. doi:10.1001/jamaneurol.2019.1424 PubMed PMID: 31157827.

64. Giannakopoulos P, Herrmann FR, Bussière T, Bouras C, Kövari E, Perl DP, et al. Tangle and neuron numbers, but not amyloid load, predict cognitive status in Alzheimer’s disease. Neurology. 2003 May 13;60(9):1495–500. doi:10.1212/01.WNL.0000063311.58879.01 PubMed PMID: 12743238.

65. Yang HS, Anzai JAU, Yau WYW, Healy BC, Román Viera AM, Maa C, et al. Plasma phosphorylated tau 217 and longitudinal trajectories of Aβ, tau, and cognition in cognitively unimpaired older adults. Nature Communications 2026 17:1. 2026 Apr 14;17(1):3188-. doi:10.1038/s41467-026-71269-3 PubMed PMID: 41980948.

66. Budd Haeberlein S, Aisen PS, Barkhof F, Chalkias S, Chen T, Cohen S, et al. Two Randomized Phase 3 Studies of Aducanumab in Early Alzheimer’s Disease. J Prev Alzheimers Dis. 2022 Apr 1;9(2):197–210. doi:10.14283/jpad.2022.30 PubMed PMID: 35542991.

67. Mintun MA, Lo AC, Evans CD, Wessels AM, Ardayfio PA, Andersen SW, et al. Donanemab in Early Alzheimer’s Disease. New England Journal of Medicine. 2021 May 6;384(18):1691–704. doi:10.1056/NEJMoa2100708

68. Sims JR, Zimmer JA, Evans CD, Lu M, Ardayfio P, Sparks J, et al. Donanemab in Early Symptomatic Alzheimer Disease: The TRAILBLAZER-ALZ 2 Randomized Clinical Trial. JAMA. 2023 Aug 8;330(6):512–27. doi:10.1001/jama.2023.13239 PubMed PMID: 37459141.

69. Dyck CH van, Swanson CJ, Aisen P, Bateman RJ, Chen C, Gee M, et al. Lecanemab in Early Alzheimer’s Disease. New England Journal of Medicine. 2023 Jan 5;22(3):142–3. doi:10.1056/nejmoa2212948 PubMed PMID: 36449413.

70. Wang Y, Mandelkow E. Tau in physiology and pathology. Nature Reviews Neuroscience 2015 17:1. 2015 Dec 3;17(1):22–35. doi:10.1038/nrn.2015.1 PubMed PMID: 26631930.

71. Mattsson-Carlgren N, Andersson E, Janelidze S, Ossenkoppele R, Insel P, Strandberg O, et al. Aβ deposition is associated with increases in soluble and phosphorylated tau that precede a positive Tau PET in Alzheimer’s disease. Sci Adv. 2020 Apr 1;6(16). doi:10.1126/sciadv.aaz2387 PubMed PMID: 32426454.

72. Bloom GS. Amyloid-β and Tau: The Trigger and Bullet in Alzheimer Disease Pathogenesis. JAMA Neurol. 2014 Apr 1;71(4):505–8. doi:10.1001/jamaneurol.2013.5847 PubMed PMID: 24493463.

73. Dujardin S, Commins C, Lathuiliere A, Beerepoot P, Fernandes AR, Kamath T V., et al. Tau molecular diversity contributes to clinical heterogeneity in Alzheimer’s disease. Nature Medicine 2020 26:8. 2020 Jun 22;26(8):1256–63. doi:10.1038/s41591-020-0938-9 PubMed PMID: 32572268.

74. Brier MR, Gordon B, Friedrichsen K, McCarthy J, Stern A, Christensen J, et al. Tau and Ab imaging, CSF measures, and cognition in Alzheimer’s disease. Sci Transl Med. 2016 May 11;8(338). doi:10.1126/scitranslmed.aaf2362 PubMed PMID: 27169802.

75. Malpetti M, Joie R La, Rabinovici GD. Tau Beats Amyloid in Predicting Brain Atrophy in Alzheimer Disease: Implications for Prognosis and Clinical Trials. Journal of Nuclear Medicine. 2022 Jun 1;63(6):830–2. doi:10.2967/JNUMED.121.263694 PubMed PMID: 35649659.

76. Arriagada P V., Growdon JH, Hedley-Whyte ET, Hyman BT. Neurofibrillary tangles but not senile plaques parallel duration and severity of Alzheimer’s disease. Neurology. 1992;42(3):631–9. doi:10.1212/wnl.42.3.631 PubMed PMID: 1549228.

77. Rosenthal SR, Guerin Camille, Weill-Halle B, Wallgren A. BCG Vaccination Against Tuberculosis. Rosenthal SR, editor. London : J. & A. Churchill; 1957.

78. World Health Organization. BCG vaccine: WHO position paper, February 2018 – Recommendations. Vaccine. 2018 Jun 7;36(24):3408–10. doi:10.1016/j.vaccine.2018.03.009 PubMed PMID: 29609965.

79. Mangtani P, Abubakar I, Ariti C, Beynon R, Pimpin L, Fine PEM, et al. Protection by BCG Vaccine Against Tuberculosis: A Systematic Review of Randomized Controlled Trials. Clinical Infectious Diseases. 2014 Feb 15;58(4):470–80. doi:10.1093/cid/cit790 PubMed PMID: 24336911.

80. Andersen P, Doherty TM. The success and failure of BCG — implications for a novel tuberculosis vaccine. Nature Reviews Microbiology 2005 3:8. 2005 Jul 11;3(8):656–62. doi:10.1038/nrmicro1211 PubMed PMID: 16012514.

81. Cernuschi T, Malvolti S, Nickels E, Friede M. Bacillus Calmette-Guérin (BCG) vaccine: A global assessment of demand and supply balance. Vaccine. 2018 Jan 25;36(4):498–506. doi:10.1016/J.VACCINE.2017.12.010 PubMed PMID: 29254839.

82. Morales A, Eidinger D, Bruce AW. Intracavitary Bacillus Calmette-Guerin in the treatment of superficial bladder tumors. J Urol. 1976 Aug;116(2):180–3. doi:10.1016/s0022-5347(17)58737-6 PubMed PMID: 820877.

83. Bannister S, Sudbury E, Villanueva P, Perrett K, Curtis N. The safety of BCG revaccination: A systematic review. Vaccine. 2021 May 12;39(20):2736–45. doi:10.1016/J.VACCINE.2020.08.016 PubMed PMID: 33810902.

84. Brausi M, Oddens J, Sylvester R, Bono A, Van De Beek C, Van Andel G, et al. Side Effects of Bacillus Calmette-Guérin (BCG) in the Treatment of Intermediate-and High-risk Ta, T1 Papillary Carcinoma of the Bladder: Results of the EORTC Genito-Urinary Cancers Group Randomised Phase 3 Study Comparing One-third Dose with Full Dose a…. Eur Urol. 2014 Jan 1;65(1):69–76. doi:10.1016/J.EURURO.2013.07.021 PubMed PMID: 23910233.

85. Cabas P, Rizzo M, Giuffrè M, Antonello RM, Trombetta C, Luzzati R, et al. BCG infection (BCGitis) following intravesical instillation for bladder cancer and time interval between treatment and presentation: A systematic review. Urologic Oncology: Seminars and Original Investigations. 2021 Feb 1;39(2):85–92. doi:10.1016/J.UROLONC.2020.11.037 PubMed PMID: 33308969.

86. Van Zijl PCM, Yadav NN. Chemical exchange saturation transfer (CEST): What is in a name and what isn’t? Magn Reson Med. 2011 Apr 1;65(4):927–48. doi:10.1002/MRM.22761;PAGE:STRING:ARTICLE/CHAPTER PubMed PMID: 21337419.

87. Milner MT, Lawrence GM, Holley CL, Bodea LG, Götz J, Burgener SS, et al. Isolation and culture of pure adult mouse microglia and astrocytes for in vitro characterization and analyses. STAR Protoc. 2022 Jun 17;3(2):101295. doi:10.1016/j.xpro.2022.101295 PubMed PMID: 35463473.

