## Supplementary Figures for "BCG vaccination mitigates tau pathology and restores cognitive function in PS19 mice"

**Title:**

**One sentence summary:**

BCG therapy prevents tauopathy in PS19 mouse model.

**Key words:**

*Mycobacterium bovis* BCG, BCG, Tauopathy, dementia, atrophy, neurodegeneration, brain, FTLD, T2-W MRI, CEST, hippocampus, RNA-sequencing, CD80, peripheral, trained immunity, glycolysis, phagocytosis.

| Institution | Subjects | % with BCG Tx | Hazard Ratio (% ↓ Alz. dementia) | *p* <0.05 | F/U (yr) | Comment |
|---|---|---|---|---|---|---|
| **1. Six retrospective human studies of BCG for dementia** | | | | | | |
| **Hebrew Unversity (Israel)** | 1371 | 64% | 0.282 (78%) | YES | 14 | |
| **MGH-Brigham (USA)** | 6467 | 52% | 0.8 (20%) Overall<br>0.7 (30%) Pt ≥ 70 yr | YES | 15 | -Decreased risk of death in patients without an earlier diagnosis of AD |
| **NYU, Montefiore (USA)** | 1290 | 25% | 0.41 (59%) | YES | 3 | -BCG effect was dose-dep.<br>-Females less responsive |
| **Hebrew Unversity (Israel)** | 12,185 | 19% | 0.72 (28%) HMO<br>0.42 (58%) Hospital | YES | 3.5–7 | -Parkinson's disease showed a reduction of 28% with BCG |
| **U. Washington Seattle (USA)** | 26,584 | 51% | 0.73 (27%) | YES | 3–4 | -NIH SEER database.<br>-BCG effect was dose-dep. |
| **Uppsala Univ., (Sweden)** | 38,934 | 17% | 0.88 (12%)<br>0.75 (25%) if age > 75 | YES | 6 | -Swedish national database<br>-Females more responsive |
| **2. Small-scale prospective study** | | | | | | |
| **Wisconsin, USA** | 49 BCG-naïve healthy humans | 100% | | YES (0.016) | 9 months | APS showed significant reduction in high-risk subjects (n = 10) after BCG doses. |

**Supplementary Figure S1.** Table explaining retrospective data, prospective data, meta-analysis demonstrating potential BCG-mediated beneficial effects against onset of dementia. (Tx- treated; F/U- Follow-up in years; APS- Amyloid probability score)

**Step 1.** Download all images from Proscia, with identical image parameters (Gamma correction, Brightness, Contrast, and Magnification).

**Step 2.** Load image in ImageJ.

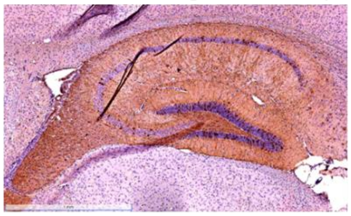

**Step 3.** Draw a line over the scale bar>Analyze>Set scale>Known distance= 1; Unit of length= mm; check “Global”> OK.

**Step 4.** Crop out scale bar.

**Step 5.** Image>color> colour deconvolution 2> Vectors: H&E DAB> OK

**Step 6.**

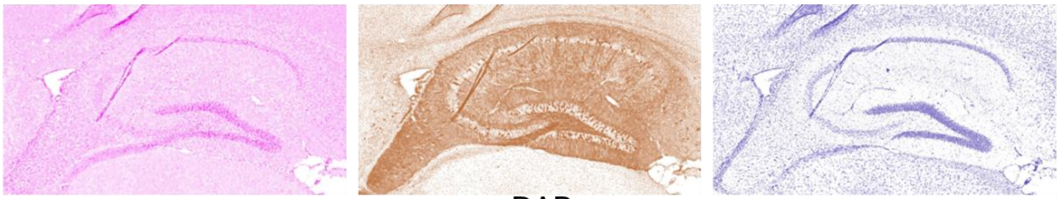

DAB

**Step 7.** Select DAB- image**.** Edit>invert.

**Step 8.** Select hippocampus area with polygon selection> Edit> Clear outside.

(Optional: Remove any obvious artifacts in the section> Select ROI with polygon selection> Edit> Clear.)

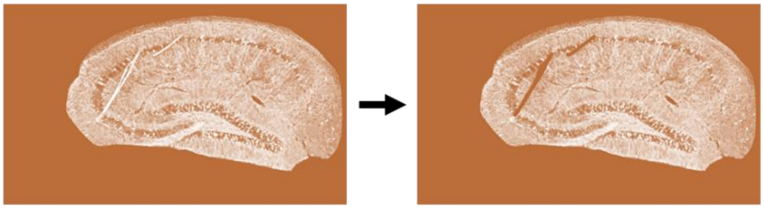

**Step 8.** Light/White shaded area should be AT8+ area. Using polygon selection, mark total hippocampus area.

**Step 9.** To remove false positives in the image, we selected strict threshold.

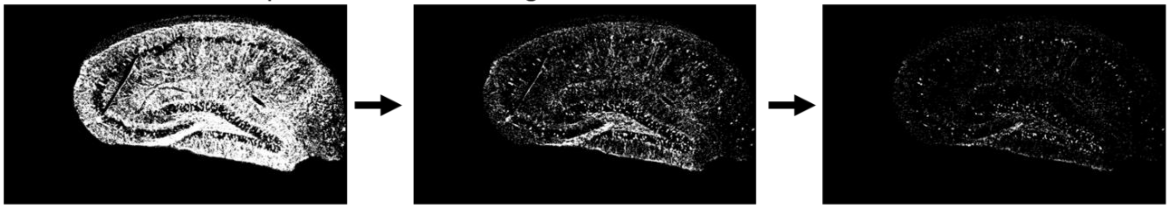

| (Threshold= 133, 255)<br>%AT8+ area= 53.65 | (Threshold=200, 255)<br>%AT8+ area= 13.02 | (Threshold=225, 255)<br>%AT8+ area= 3.27 |
|---|---|---|

**Step 10.** Analyze>Measure> Record “%Area”.

**Supplementary Figure S2. ImageJ-analysis pipeline used to quantify immunohistochemistry-positive area in the brain sections, as shown in Fig. 1.** For each mouse brain sample, IHC was done in three planes (10 µm thick sections) for all the antibodies.

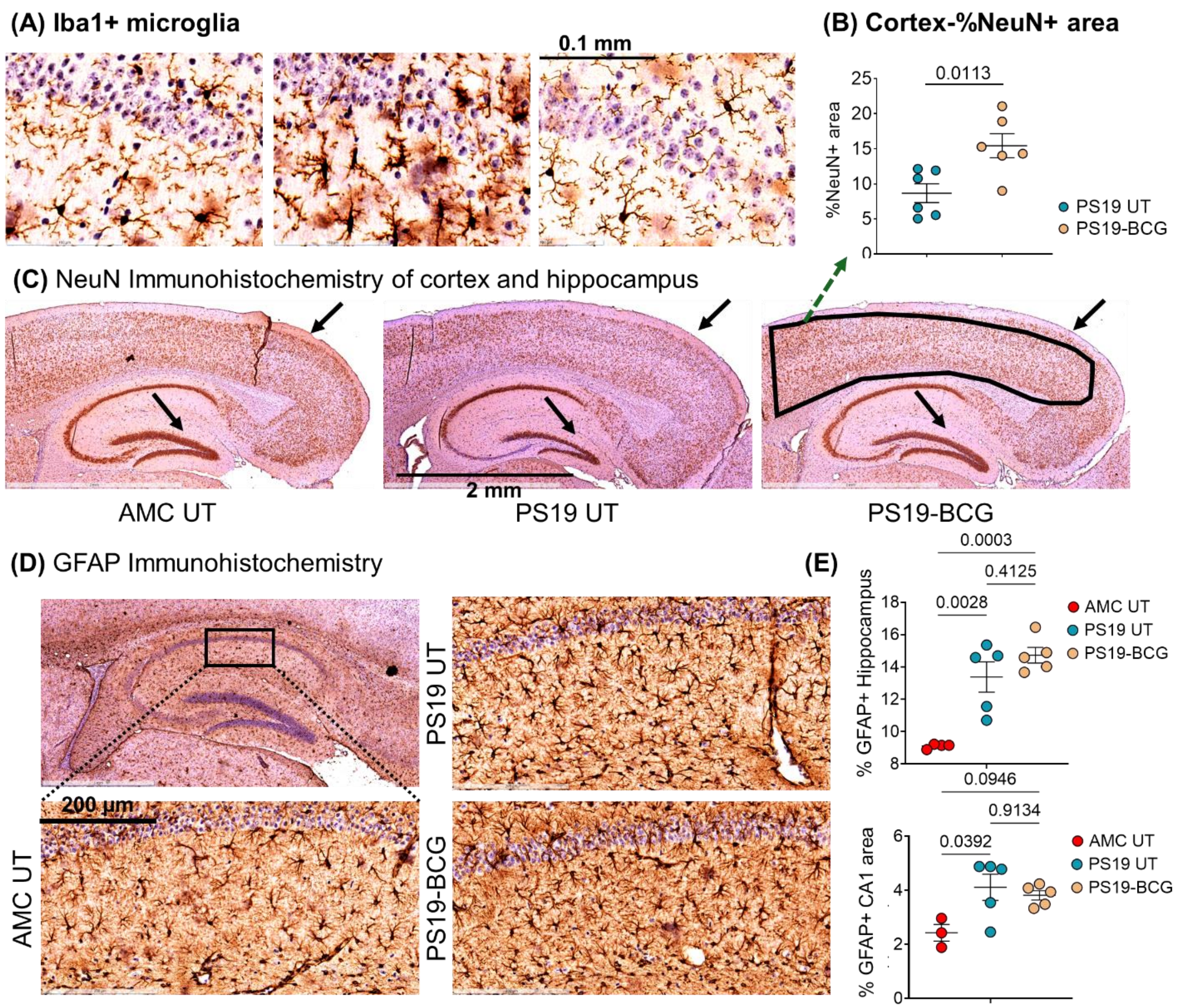


**Supplementary Figure S3.** ***M. bovis*** **BCG treatment ameliorates neurodegeneration, and microgliosis in PS19 mice, but did not reduce astrocyte activation levels. (A)** Representative image of Iba1-IHC area in the hippocampus-CA1 region of AMC UT, Sham (PBS) or BCG treated PS19 mice, showing restoration of homeostatic, ramified microglia in the BCG-treated PS19 mice. **(B, C)** %NeuN+ area in the cortex region (somatosensory and visual area) of AMC UT, Sham (PBS) or BCG treated PS19 mice, as determined by ImageJ analysis. (n=6/group) Black arrow indicates differences in NeuN-IHC staining. **(D, E)** %GFAP+ area in the hippocampus and hippocampus-CA1 region of AMC UT, Sham (PBS) or BCG treated PS19 mice, as determined by ImageJ analysis. (n=6/group). The data are means ± SEM, and representative of two independent experiments. Each data point represents a mouse. Statistical analysis was calculated by unpaired two-tailed Student's t-test (B) or one-way ANOVA with Tukey's multiple comparisons test (E). ($p$>0.05: ns, $p$<0.01: **, $p$<0.001: ***, $p$<0.0001: ****)

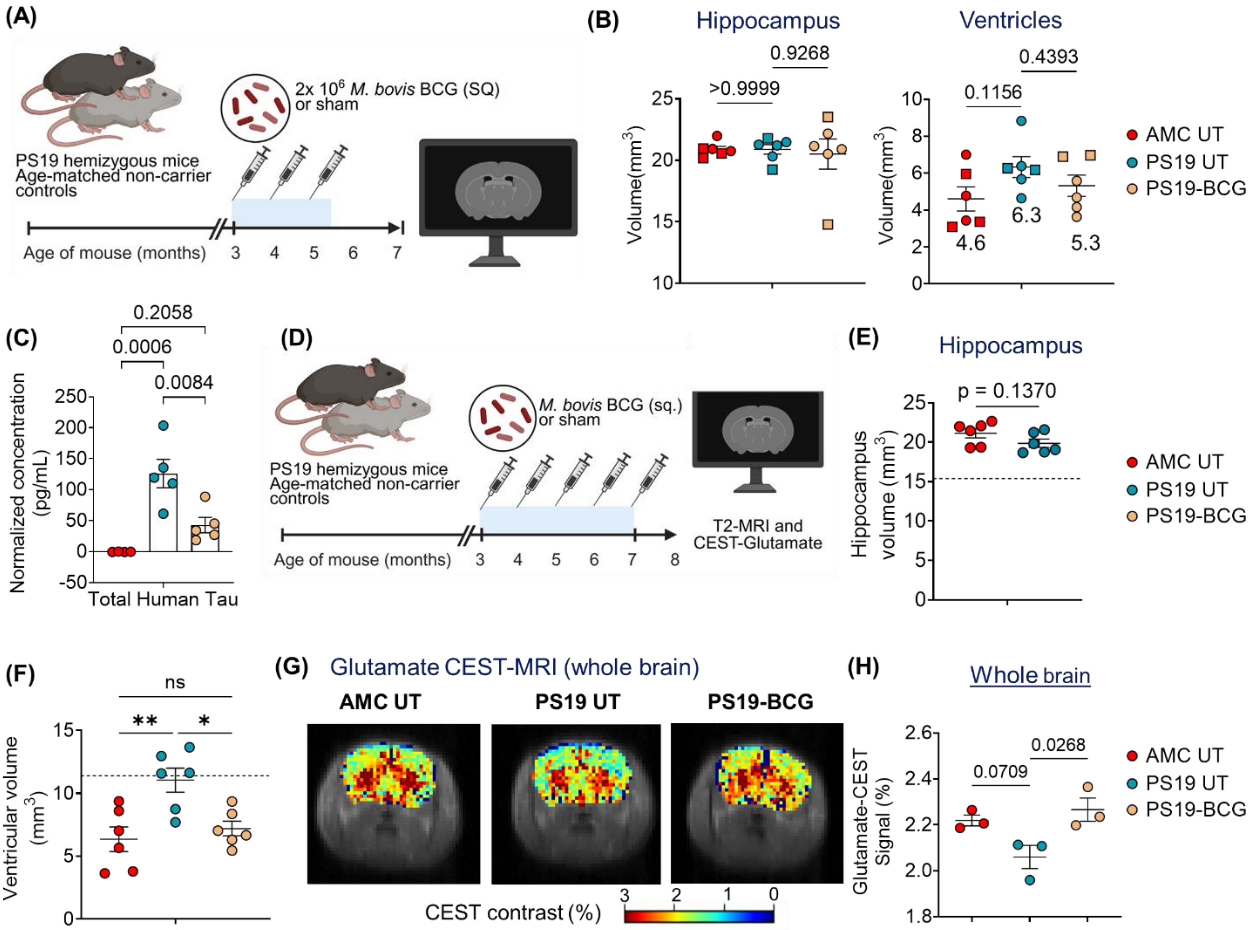


**Supplementary Figure S4. *M. bovis* BCG treatment prevented ventricular dilatation in the brain of PS19 males in the cohort 2. (A)** Schematic of the experiment-Cohort 1. **(B)** Quantitative analysis of the hippocampus and lateral ventricle volume. (n=6/group; 2-3 males (squares), 3-4 females(circles)) The data are means ± SEM. Each data point represents a mouse. Statistical analysis was calculated by one-way ANOVA with Sidak's multiple comparisons test. Mean ventricular volume are mentioned on the figure. **(C)** Total human tau protein levels detected in plasma from 6-7 months old PS19 mice. n= 5 mice/group. **(D)** Schematic of the experiment-Cohort 2. **(E, F)** Quantitative analysis of the hippocampus and lateral ventricle volume of PS19 males. n=6/group. **(G, H)** Representative image of glutamate-weighted chemical exchange saturation transfer (CEST-MRI) maps of the brain. (Cohort 1; n=3 mice/group), and quantification. The data are means ± SEM, and representative of two independent experiments. Each data point represents a mouse. Statistical analysis was calculated by unpaired two-tailed Student's t test (Hippocampus in **E**) and one-way ANOVA with Sidak's multiple comparisons test (**C, F, H**). (*p*>0.05: ns, *p*<0.01: **, *p*<0.001: ***, *p*<0.0001: ****)

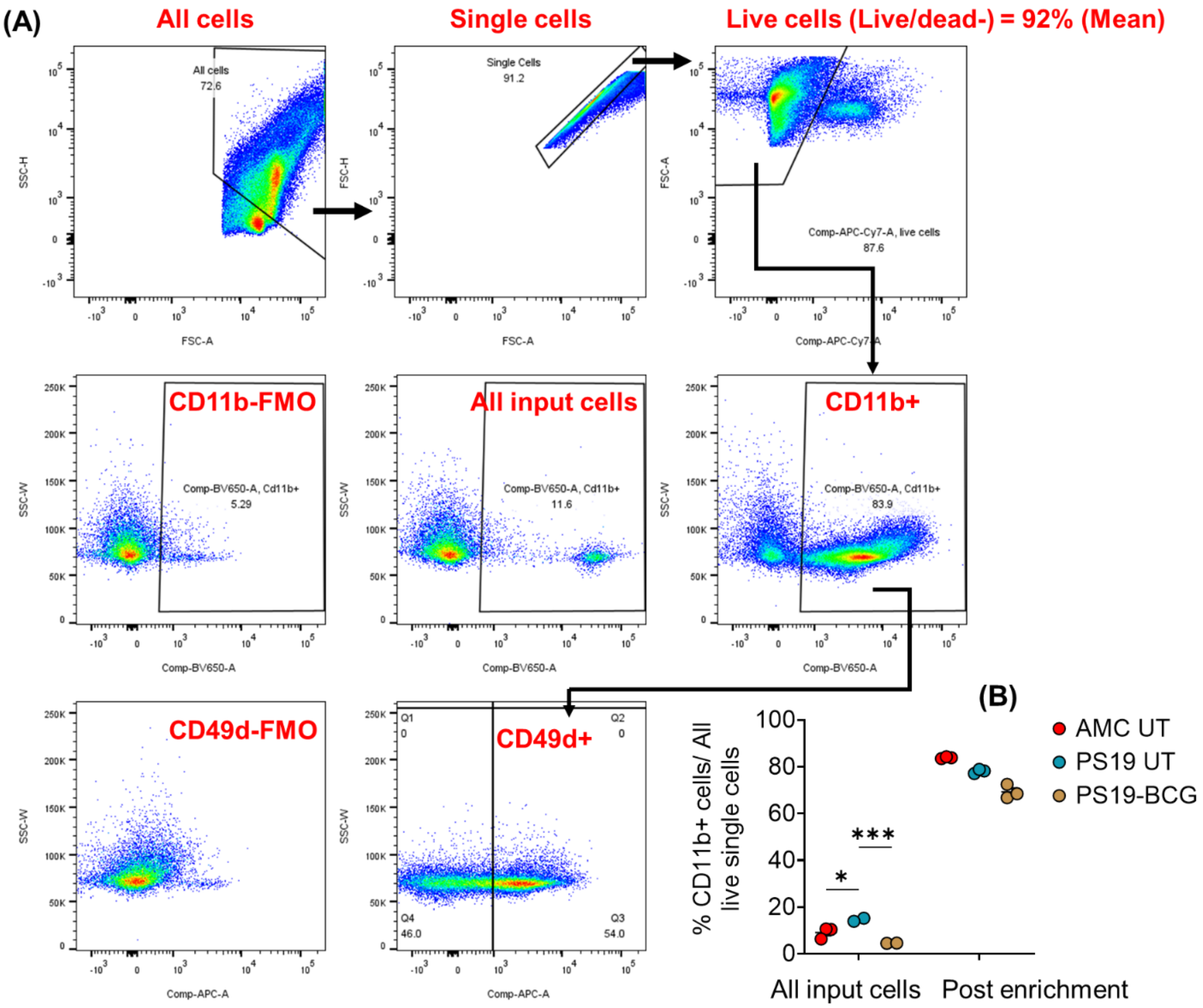


**Supplementary Figure S5. Immunophenotyping of CD11b+ cells from PS19 mouse brain. (A)** Representative flow cytometry dot plots and arrows depicting gating strategy and pre-gated cell populations. Gating for all the positively-stained events were established based on respective Fluorescence minus one (FMO) control. The mean viability of the CD11b+ cells derived from magnetic-bead based enrichment was 92%. **(B)** Higher CD11b+ cells out of all live single cells were detected in sham-treated PS19 brains, before the enrichment by magnetic beads (input cells). (n=6 mice/group; samples from 2 mice were pooled together) The data are means ± SEM, and representative of two independent experiments. Each data point represents a mouse. Statistical analysis was calculated by two-way ANOVA with Tukey's multiple comparisons test. ($p$>0.05: ns, $p$<0.01: **, $p$<0.001: ***)
